# Combining molecular dynamics simulations and scoring method to computationally model ubiquitylated linker histones in chromatosomes

**DOI:** 10.1101/2022.09.01.506295

**Authors:** Kevin Sawade, Christine Peter, Andreas Marx, Oleksandra Kukharenko

**Affiliations:** Department of Chemistry, University of Konstanz, Universitätsstraße 10, 78457 Konstanz, Germany; Max-Planck Institute for Polymer Research, Ackermannweg 10, 55128 Mainz, Germany

## Abstract

The chromatin in eukaryotic cells plays a fundamental role in all processes during a cell’s life cycle. This nucleoprotein is normally tightly packed but needs to be unpacked for expression and division. The linker histones are critical for such packaging processes and while most experimental and simulation works recognize their crucial importance, the focus is nearly always set on the nucleosome as the basic chromatin building block. Linker histones can undergo several modifications, but only few studies on their ubiquitylation have been conducted. Mono-ubiquitylated linker histones (HUb), while poorly understood, are expected to influence DNA compaction. The size of ubiquitin and the globular domain of the linker histone are comparable and one would expect an increased disorder upon ubiquitylation of the linker histone. However, the formation of higher order chromatin is not hindered and ubiquitylation of the linker histone may even promote gene expression. Structural data on chromatosomes is rare and HUb has never been modeled in a chromatosome so far. Descriptions of the chromatin complex with HUb would greatly benefit from computational structural data. In this study we generate molecular dynamics simulation data for six differently linked HUb variants with the help of a sampling scheme tailored to drive the exploration of phase space. We identify conformational sub-states of the six HUb variants using the sketch-map algorithm for dimensionality reduction and iterative HDBSCAN for clustering on the excessively sampled, shallow free energy landscapes. We present a highly efficient geometric scoring method to identify sub-states of HUb that fit into the nucleosome. We predict HUb conformations inside a nucleosome using on-dyad and off-dyad chromatosome structures as reference and show that unbiased simulations of HUb produce significantly more fitting than non-fitting HUb conformations. A tetranucleosome array is used to show that ubiquitylation can even occur in chromatin without too much steric clashes.

**Author summary:** In eukaryotic cells the linker histones play a crucial role in the formation of higher order nucleoprotein complex of DNA, especially for the arrangement of the nucleosomes. Histones can undergo several modifications, but modification of a linker histone with a single udiquitin (mono-ubiquitylation) remains one of the least understood epigenetic modifications. One reason is the inaccessibility of homogeneously modified linker histones for experimental methods, which are crucial for distinct studies. We combine molecular dynamics simulations with machine learning-based approaches to study the influence of mono-ubiquitylation in linker histones on DNA interaction and their ability to form higher order chromatin structures. We were able to determine the probable states in six differently linked histone-ubiquitin complexes via accelerating classical molecular dynamics simulations and using advanced state characterization techniques. As it is computationally unfeasible to simulate the whole chromatosome with different modified histones we developed efficient geometric scoring technique to select biologically relevant structures of all six mono-ubiquitylated linker histone that can bound to nucleosome.

## Introduction

Chromatin is the nucleoprotein complex in cells in which DNA is packed. Besides the DNA it is composed of various histone and non-histone proteins. [1] The basic building block of chromatin is the nucleosome, a complex which is formed when ≈ 147–167 bp of DNA wrap around an octameric protein complex of four pairs of core histones. [2] In chromatin, the nucleosome is accompanied by a linker histone and this complex is then called chromatosome. The linker histone interacts with the DNA linkers which connect the chromatosomes and thus directly influences higher packing of the chromatin [3], such as the formation and structure of the 30 nm chromatin fiber. The nature of linker histone binding modes and chromatin packing is a topic of ongoing research. [4–10]

The binding affinity of the linker histone to the chromatin can be affected by the histone’s post-translational modifications, such as methylation, acetylation, phosphorylation, etc. [11] One such post-translational modification is the formation of an isopeptide bond between a substrate protein’s lysine residue and the C-terminus of the protein ubiquitin (Ub) (termed as “ubiquitylation” or “ubiquitination”). Ubiquitylation is known to occur at the core histones, where it can regulate, among others, gene silencing, [12] chromatosome dynamics, and ultimately DNA accessibility [13]. Later ubiquitylation was discovered to also occur on the linker histone (Fig 1). [14–16] Furthermore, polyubiquitylation of the linker histone has been identified as an intermediate in the repair of double strand DNA break. [17, 18] The chromatin system has also been studied in silico in various resolutions ranging from highly abstracted models where the nucleosome comprises a rigid body to multiscale and all-atom simulations of the nucleosome core structure. [19–23] However, most of the studies exclude the linker histone from their models, despite acknowledging the importance of this protein in the formation mechanism of higher order chromatin. The position of the linker histone in the “pocket” of linker and core DNA, i.e. the binding modes, has been subject of discussion but most probably varies between a symmetric on-dyad mode and an asymmetric off-dyad mode. [10, 24–27] Recently, all-atom simulations have been conducted to further investigate the nucleosome/chromatosome system. [22, 23, 28] Yet, ubiquitylation of the linker histone remains underrepresented.

**Figure 1.**
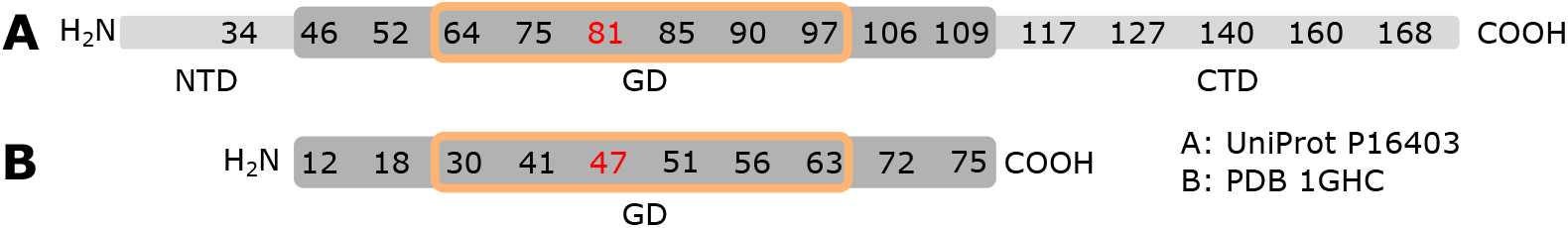
Schematic sequences of wild type human linker histone H1.2 (**A**) and the globular domain (dark grey) of chicken histone H1 (PDB 1GHC [29] **B**). The numbers represent aligned lysine residues that have been found to undergo ubiquitylation. [14–16] The red numbers indicate lysines considered in this study that have not been found to be ubiquitylated in nature (K47 in 1GHC and K81 in human H1.2). Lysines within the orange outline are investigated in this study. Abbreviations: NTD (N-terminal domain), GD (globular domain), CTD (C-terminal domain).

First investigations assessing the influence of ubiquitylated linker histone (HUb) on the chromatosome complex suggested a trend towards a compaction of chromatosome arrays upon ubiquitylation. [30] However, some studies have shown that ubiquitylation of histones and especially K30 ubiquitylated linker histone (K64 in human H1.2, see Fig 1 for comparison of lysines’ numbering) prevent compaction, relax the structure, and promote transcription. [31, 32] Given the geometry of the system it seems astounding that Ub can form a covalent bond to the similar sized globular domain of the linker histone while still allowing a functional and ordered complex to form. Even more so, when an array of multiple chromatosomes is considered. [33] *Structural data on ubiquitylated linker histones is still not available*.

As many cellular processes depend on chromatin accessibility, static or dynamic structural data of HUb and HUb inside a chromatosome would greatly benefit from further research. Structural data on the chromatosome is sparse: the protein database lists 8 such structures, as of this study. These chromatosomes mainly vary in the length of the linker DNA. All of them exhibit the on-dyad binding mode. We decided to use the three chromatosome structures obtained via X-ray crystallography as references (first three entries in Table 1). All of these structures are synthetic biological constructs reconstituted from core histones and the Widom 601 DNA sequence, which has been specifically tailored to increase binding to the core histones and thus has come under criticism. [34, 35] No structures let alone dynamic data of ubiquitylated linker histone have yet been published. Attempts to crystallize HUb in a chromatosome complex have not been fruitful so far. [36] In this study, we rely on the structural data summarized in Table 1 as basis for all further modeling. The structure in off-dyad binding mode obtained by molecular docking was kindly provided by Dr. Yawen Bai. [10] A notable example is the tetranucleosome array PDB 1ZBB crystallized by Schalch et al. [4] which we will use to bridge the gap between single nucleosome/chromatosome structures and higher order chromatin.

**Table 1.**
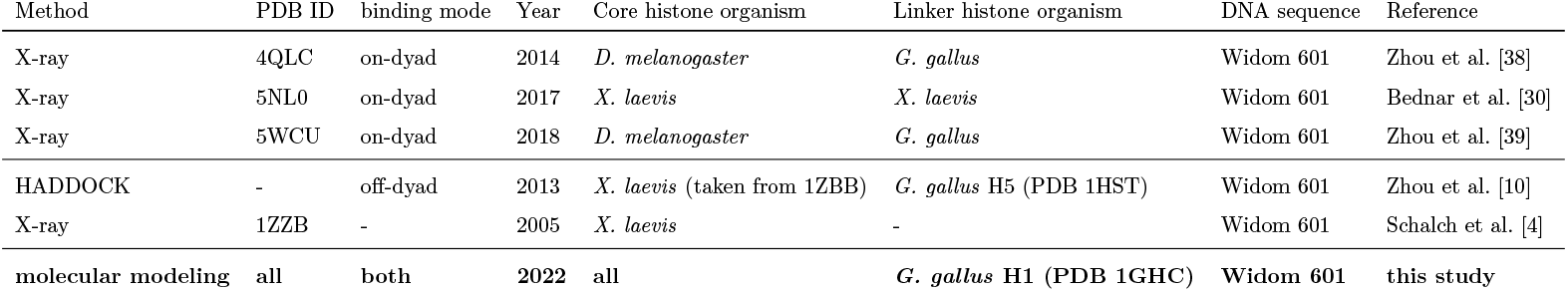
Five chromatosome structures used as reference in this study: PDBs 4QLC, 5NL0, and 5WCU (rows 1-3) exhibit an on-dyad binding mode of the linker histone. The reference structure of the off-dyad binding mode (row 5) was obtained via HADDOCK docking using a nucleosome from 1ZBB and PDB 1HST as the histone. [10, 37]. PDB 1ZBB is used as a reference for a an array of multiple nucleosomes. The last row represents the present study. We use PDB 1GHC as the linker histone (also see Fig 1 for a sequence alignment of the linker histones).

The X-ray and HADDOCK methods provide valuable first steps for structural data on the chromatosome. However, they strive to find a single structure of an inherently dynamic system. In this paper we aim for the combination of molecular dynamics simulations with machine learning techniques to explore the conformational landscape of ubiquitylated linker histone. We offer possible conformations of HUb in chromatosomes and chromatosome arrays to reveal a novel view into DNA regulation by ubiquitylation. We generate an extensive library of conformations of differently ubiquitylated linker histones using molecular dynamics (MD) simulations guided by our own enhanced sampling algorithm [40]. We reduce the data size by excluding statistically insignificant/physically improbable structures through clustering in a dimensionally-reduced space. We present a particularly efficient geometric scoring method for this system which is then used to analyze if and how the ubiquitylated linker histone fits into the chromatosome - and if this is dependent on the ubiquitination site. Last but not least we use the same methods on a tetranucleosome array to bridge the gap between isolated chromatosomes and chromatin fibers. The workflow employed in this study is visualized in Fig 2. It can be summarized into three main steps: 1) exploration of conformational phase space; 2) identification of representative conformational states; 3) scoring of HUb in on-dyad and off-dyad mono chromatosomes, and a tetranucleosome array.

**Figure 2.**
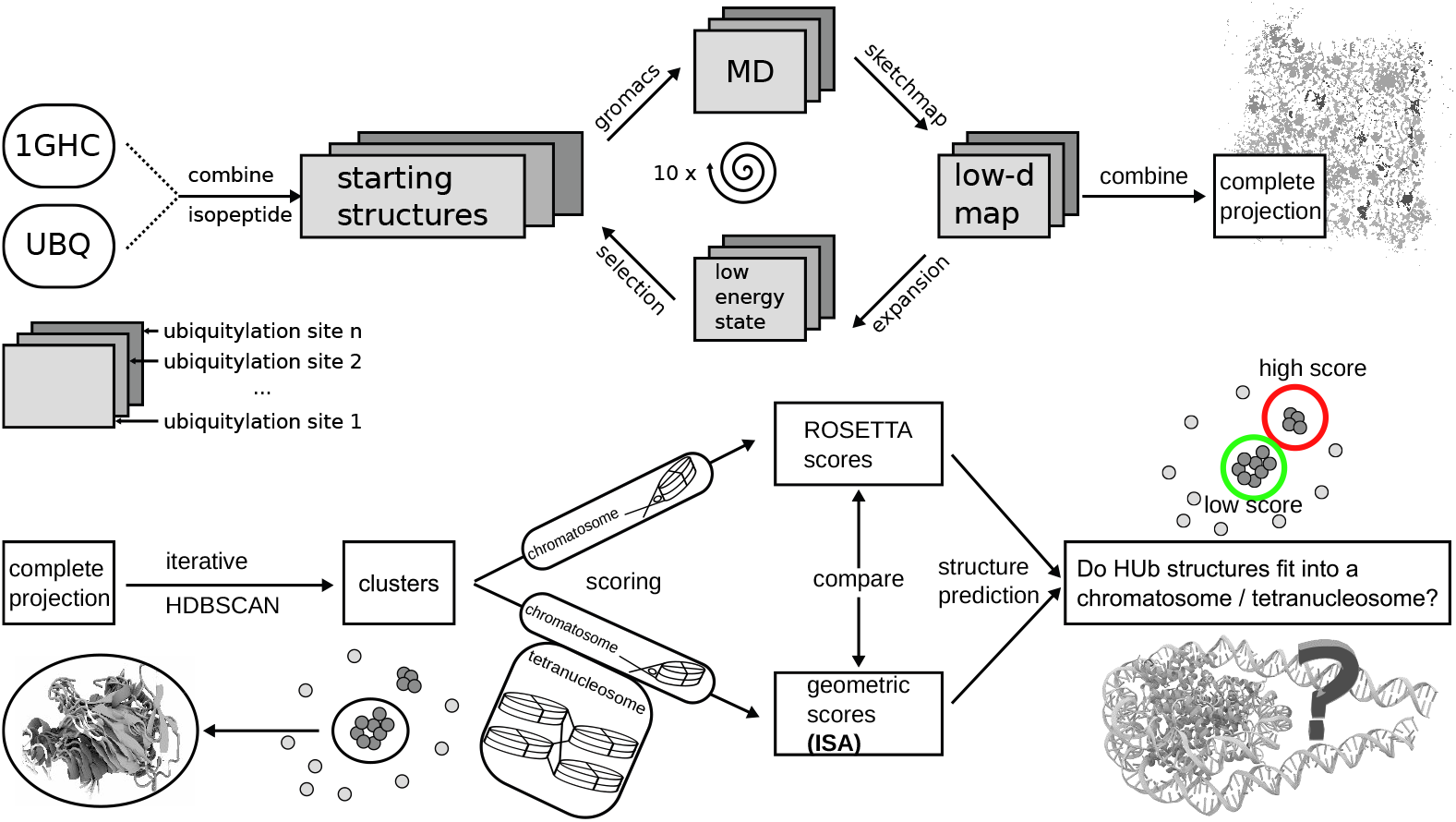
Illustration of data-generation (top half) and analysis (lower half) workflow to identify possible structures of ubiquitylated linker histone in a chromatosome. Data was produced by MD simulations driven by an expansion scheme which selects new starting structures from sparsely populated regions of phase space. The aggregated MD data is projected into a combined low-dimensional space and clustered iteratively. Clusters (i.e. characteristic conformational states) are then fitted into different chromatosome structures using a new geometric scoring method.

## Results

We make general assessments of the conformational properties of the six HUb variants and propose the identified clusters as structurally stable sub-states to be used as input for the interpenetration and scoring algorithm presented in Materials and Methods. Doing so, we are able to screen through all simulation frames and find HUb conformations that fit into the DNA “pocket” of a nucleosome and a tetranucleosome array.

### Structural diversity of six differently ubiqutylated histones

The complete set of 1180 simulations (≈ 35 μs of aggregated simulated time) of the six HUb variants was projected into the same low-dimensional space. In this projection the closeness of any two points is related to the structural similarity of the respective protein’s conformations. Patches with high density contain similar structures of very little RMSD variance, whereas solitary points represent statistically insignificant structures which can be – especially after the exhaustive sampling conducted here – viewed as transition states. RMSDs between structures belonging to high-density basins are proportional to their respective distance in sketch-map space, however, they are not linearly related (Fig 3 (B) and (E)). Using this method we could identify large regions of unique structures for every ubiquitylation variant and small intersections of these regions, where the conformational space of two or more variants overlap (Fig 3 (A)). The starting sidechain dihedral angle χ_3_ is the second most prominent feature (Fig 3 (D)). These map regions can not be taken as statistical weights for certain conformations as for that a notion of density is required (Fig 3 (C)). After the removal of high density regions (S1 Text) the density can be compared with the similar di-Ubiquitin protein (diUb). The density map is more shallow than similar diUbs. [41] This is due to the histone’s conformational space which is greater, when compared to ubiquitin (i.e. the histone is more flexible) and the expansion scheme driving the simulation away from meta-stable, local-minima conformations. We additionally questioned the completeness of the conducted sampling and stability of obtained meta-stable states, but assessing ergodicity is out of the scope of this work.

**Figure 3.**
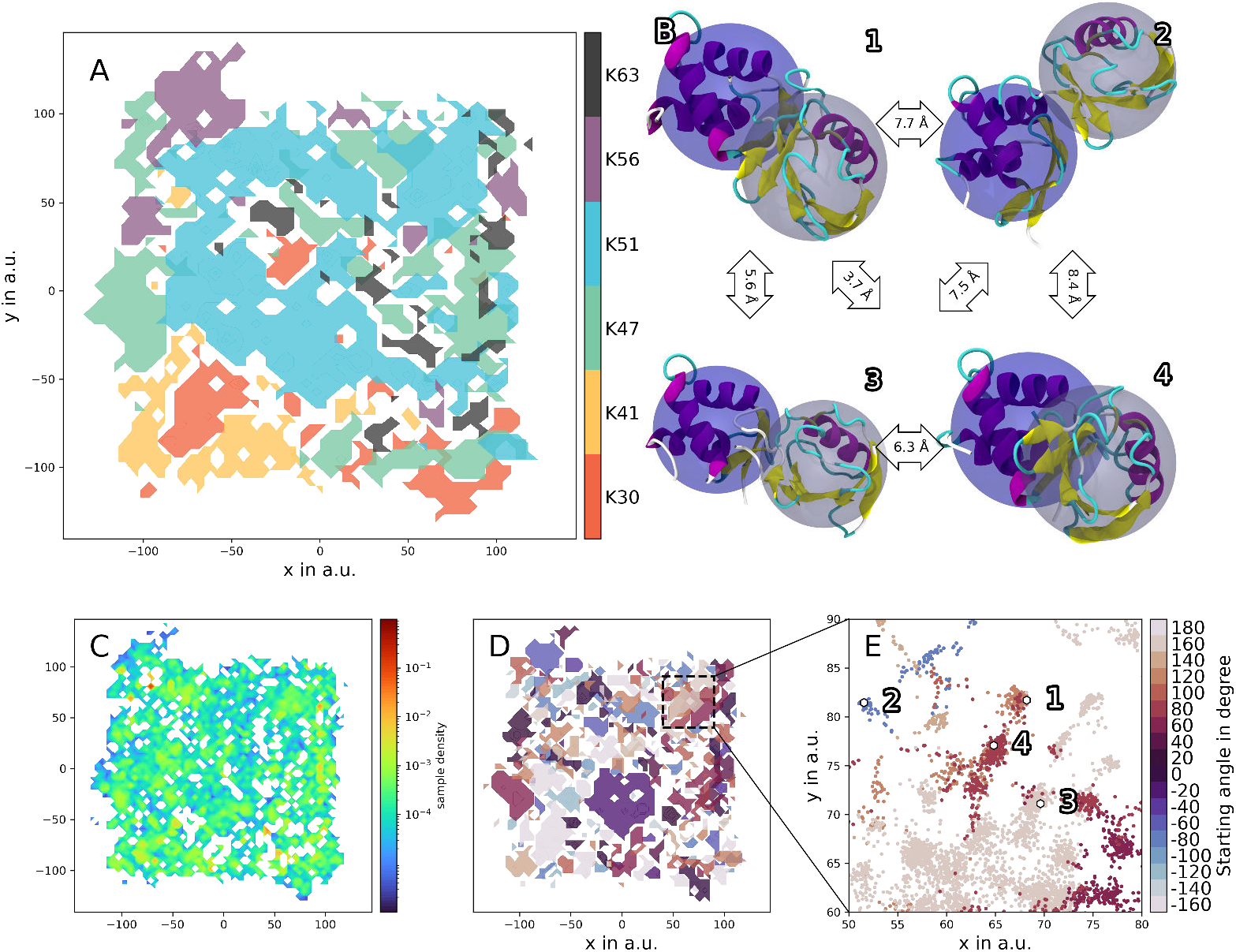
Sketch-map projections of all conducted simulations colored by various parameters and exemplary structures. A contour plot with transparent patches colored according to ubiquitylation site (A) shows that the most distinct feature of the six HUb variants is the ubiquitylation site. A density map of the combined projections (C) shows that the low-dimensional representation is mostly shallow (blue) with only small regions of higher density (red). Coloring the combined projection according to the starting χ_3_ angles ((D) and (E)) shows a finer structure of the projection. Four of the basins in (E) have been chosen and their RMSD centroids are visualized with cartoon representation and colored according to secondary structure features (B). The blue transparent sphere is focused on the histone subunit, the grey one on the Ub subunit. Closeness in sketch-map space is related to structural similarity. In (A) two regions of high density and high RMSD variance were excluded, to not bias the density towards these regions. Subfigure (C) was rendered after removing the high-denisty, high-RMSD variance region (S1 Text).

After creating the exhaustive structural library for all six HUb variants we can address more interesting questions on: how many meta-stable states does HUb assume and which of these states can bind to a nucleosome.

### Identification of characteristic conformational states of HUb

The visual inspection of the 2D sketch-map projections already reveals the enormity of the conformational landscape of the six variants. Each point in sketch-map space represents one conformation sampled by an unbiased MD simulation. High density regions represent statistically significant sub-states of the variant’s combined phase space. The task of identifying these states can be reduced to a grouping of similar features – clustering.

Considering size and properties of the given data there are some requirements to a clustering method to be able to define meta-stable states in the 2D sketch-map projection: 1) handle vast amount of points; 2) work without recurring a predefined number of clusters; 3) distinguish between dense basins of any size and shape (meta-stable configurations) and low density background (transitions states).

We found the hierarchical density-based clustering algorithm HDBSCAN with iterative application particularly well suited for this task. HDBSCAN transforms the space before building a single linkage tree thus pushing “noise” (sparse regions) further away from the dense data. We applied this algorithm iteratively, first to exclude high density regions with high RMSD variance (artifacts of the projection algorithm, see S1 Text), and then to gradually define clusters proceeding from large to small by decrementing the hyperparameter (more details on the algorithm can be found in Materials and Methods: Identifying characteristic conformational states of HUb).

With this iterative clustering approach we extracted 707 clusters which represent ≈ 30% of the variants’s conformational space (Fig 4). Compared to just observing the 2D projections of the phase spaces of differently linked HUb, clustering allowed us to better analyse the joint phase space of HUb’s. Out of the 707 clusters only 64 (4% of all clustered points, 1% of all points) were sampled by a single trajectory. This points towards a convergence of the conformational space sampled by the MD simulations. A fifth of all clustered points (keep in mind that clusters can have different populations) are part of clusters that contain structures from multiple topological different variants. This implies meta-stable HUb states, which are not specific to the linker position, and this can be a hint to the question, why no significant difference can be seen in the experiments for different ubiquitylation sites. [36] The presence of clusters with differently ubiquitylated variants also confirms the advantage of using our expansion scheme compared to straightforward MD, as applying sketch-map and HDBSCAN to the initial simulations (S2 Fig) resulted in clusters exclusively built from single trajectories.

**Figure 4.**
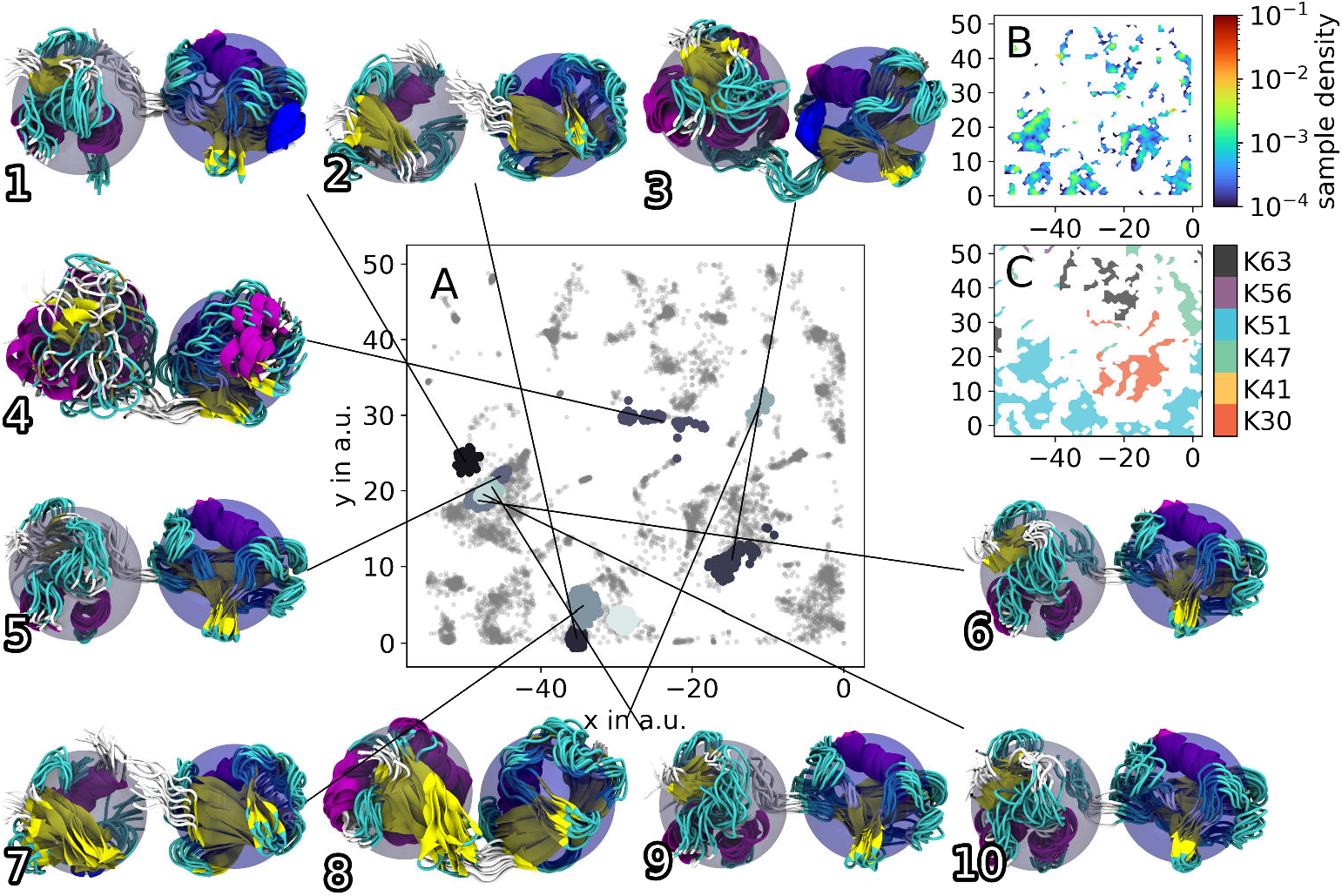
Applying an iterative HDBSCAN clustering, a total of 707 clusters were obtained from the complete sketch-map projection. A selection of a few clusters from a section of the joint projection (compare Fig 3 A, C, D for overview) is visualized as colored patches, whereas the outliers are grey and slightly transparent (A). Structure renders of these clusters (blue sphere centered on H1 subunit, grey sphere centered on Ub subunit) show that a satisfying degree of structural cohesion could be achieved. Cluster 4 represents an edge case of a cluster that was still considered within the boundaries of the iterative HDBSCAN algorithm. The non-cohesiveness of this cluster can be seen in the render and the low-dimensional projection. Subfigures (B) and (C) show the sample density and the dominant ubiquitylation site of the region in (A), respectively.

### Scoring HUb conformations into nucleosome structures

#### Scoring single structures into nucleosomes

The interpenetration and scoring algorithm (ISA) allows a fast scoring of all simulated HUb conformations (see Materials and Methods: Fitting HUb into the chromatosome). As a first introduction to the ISA scores we will use K30Ub and place it into the 5NL0 nucleosome. The sketch-map projection can now be colored by the score, revealing distinct regions of higher and lower scores (Fig 5 (A)). The lowest ISA score observed for placing K30Ub into 5NL0 is 8.4 and the highest is 406.2. The previously identified clusters represent (meta-)stable sub-states on the protein’s folding pathway and as such are good candidates to identify states of lower scores which are also likely to occur in real-life systems.

**Figure 5.**
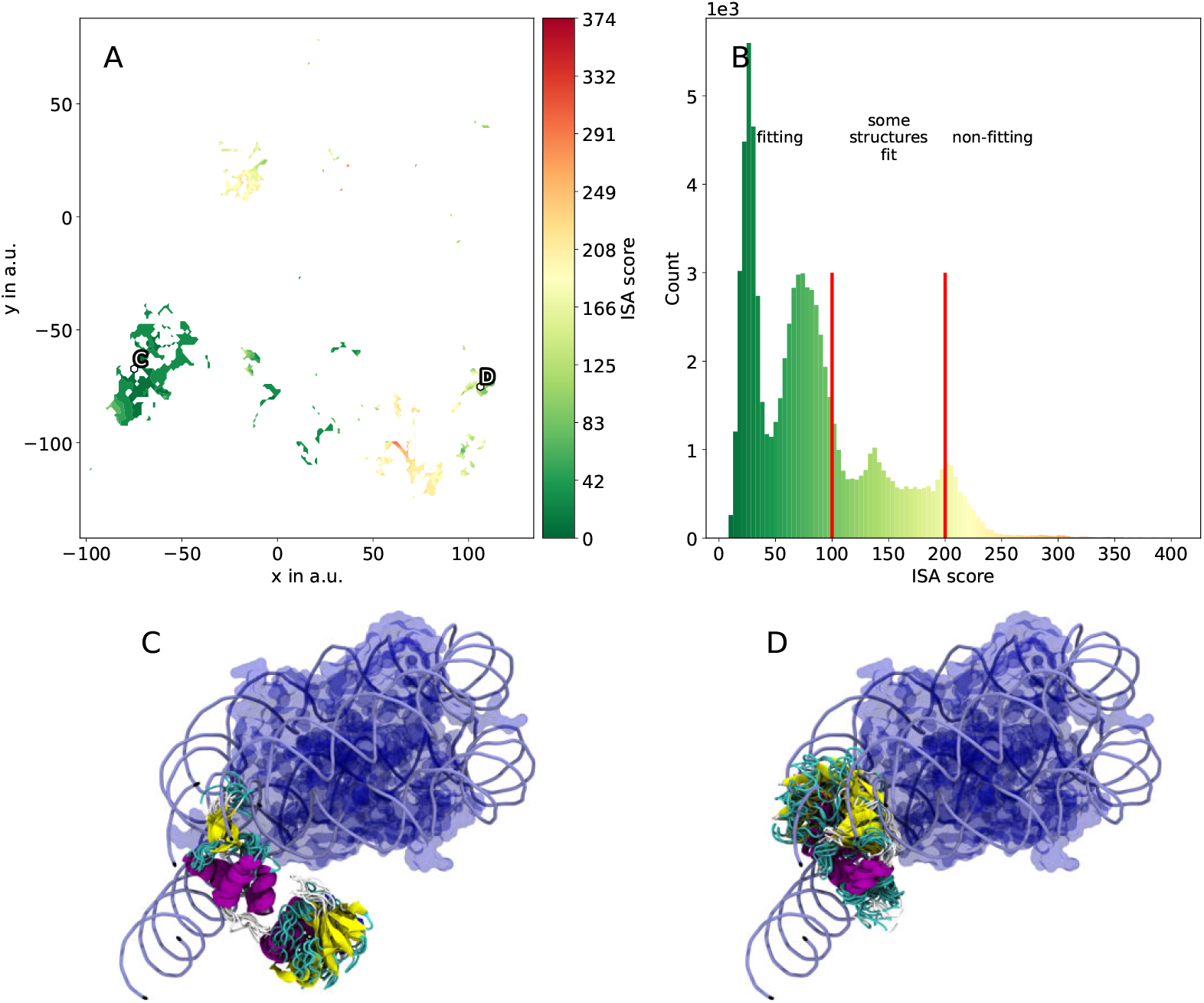
Results of the interpenetration and scoring algorithm scoring and example structures for scoring K30HUb into 5NL0. The sketch-map projection (A) colored according to ISA score exhibits distinct regions with fitting (green, low score) and non-fitting (red, high score) conformations. The histogram of scores has a skew to lower scores (B). In (C) and (D) exemplary HUb clusters are rendered within the nucleosome of 5NL0. Both (C) and (D) visualize the DNA as tubes and the core histones as a transparent blue surface. The HUb structure bundles are rendered using their secondary structure and colored accordingly. In (C) the K30HUb cluster with the lowest ISA score is shown. Using this cluster the HUb chromatosome complex could easily be used as input for further simulations. In (D) a cluster with an ISA score of 150 is shown. This cluster can be taken as an example of a high-score cluster, that does only slightly intersects with the DNA linkers.

We used 4 different chromatosome structures as receptors (rows 1-3 on-dyad H1 binding mode, row 4 off-dyad binding mode in Table 1). Thus, the resulting scores can not only be split up into the different ubiquitylation sites (Fig 6 (C)) but also into the different receptor nucleosomes (Fig 6 (E)). Although the number of atoms varies for these nucleosome structures we still can assume that low and high scores can be compared within the margin of error. A score of 0 is achievable when no intersections occurs at all. ISA scores ranging from 0 to 473 were observed. We chose an ISA score of 100 (the 75th percentile of all scores lies at 89.9, which after visual inspection of edge cases was deemed a good choice (S9 Fig) and conformations with scores lower than 100 were designated as “fitting”. Structures in between scores of 100 and 150 typically intersect with the DNA linkers (e.g. Fig 5 (D)) or the nucleosomal core. If an intersection with the DNA linker occurs the structure could potentially still be used for MD simulations after energy minimization, which is not pursued here. Protein conformations with scores of 150 and above are designated as not fitting.

**Figure 6.**
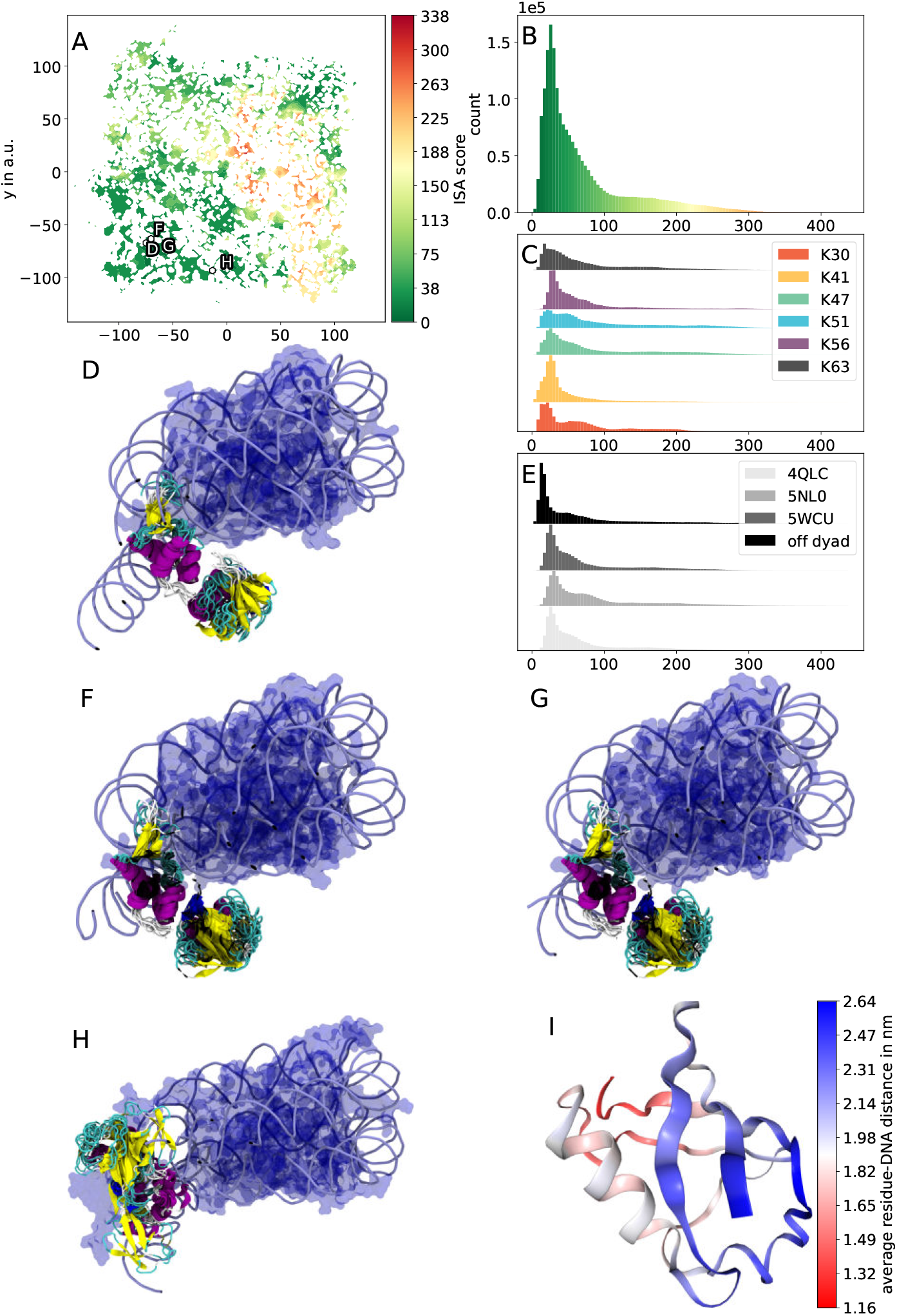
Scores and renders of the six HUb variants in the four chromatosome structures. In (A) the sketch-map projection is colored according to the ISA score where a specific region exhibits high scores and contains non-fitting HUb conformations. The distribution of scores can be seen in (B), (C), and (E). The clusters with the lowest scores of the parent chromatosomes 5NL0, 5WCU, 4QLC, and the off-dyad structure are shown in (D-H), respectively. Coloration of renders is in line with Fig 5. Note, that the lowest scoring cluster for 5WCU (F) and 4QLC (G) is the same cluster. Both chromatosomes are structurally very similar, missing the DNA linkers that are present in 5NL0. Also note, how the Ub subunit of the lowest scoring cluster fitted into the off-dyad chromatosome points into a different direction. The off-dyad structure in (H) contains the tail domains of the core histones. In (I), the crystal structure of PDB 1UBQ was colored according to the per-residue distance to DNA after placing HUb into 5NL0. The possible values range from red (closest) to blue (farthest).

The distribution of scores is skewed to lower scores. Although the HUb proteins were simulated in water without any nucleic acid present, the overall majority of sampled structures fits into the nucleosome (Fig 5 (B)). This coincides with studies observing reconstituted HUb chromatosomes to still form orderly complexes and not break the complex completely. [31] The chromatosome with off-dyad binding linker histone exhibits greater deviations because its core histones contain the flexible tails, that the other chromatosomes lack. The superposed conformations of the ubiquitylated linker histones can overlap with these regions. However, due to the flexibility of these tails [42] we expect the tails to be able to accommodate for the ubiquitin subunit and possibly interact with it in a dynamic system.

#### Scoring clusters into nucleosomes

When fitting clusters (i.e. bundles of 100 to 1000 HUb conformations) into the chromatosomes, the ubiquitin subunit does often fan out. Despite this structural variance and the entailing increase in the conformational demand of the Ub subunits, the majority of the obtained clusters did fit surprisingly well into the chromatosomes. All clusters do exhibit a non-zero mean ISA score, with the best fitting cluster (Fig 5 (C)) having a mean ISA score of 17. Using the four chromatosomes from Table 1 we found that for 4QLC and 5WCU the minimal ISA score is achieved by the same cluster (Fig 6). Interestingly, the HUb structure bundles of all on-dyad chromatosomes exhibit similar spatial positioning. Here, the Ub subunit of the lowest scoring cluster is situated “below” the nucleosomal core. In contrast, the off-dyad nucleosome the DNA linkers exit the nucleosomal core at other angles than in the on-dyad nucleosomes. Furthermore, the linker histone in the off-dyad nucleosome is slightly shifted and rotated. This leads to the lowest scoring clusters for the off-dyad nucleosome being distinctively different from the on-dyad nucleosomes. Here, the Ub subunit points upward. Using the scores of clusters the question *whether ubiquitylated linker histone can fit into the chromatosome* could be answered. As most of the clusters exhibit a lower score we assume that HUb has a tendency to assume conformations that fit into the DNA pocket of these mono-nucleosomes. However, biological systems of chromatin are comprised of many nucleosomal building blocks. Thus, investigating the positioning of HUb in an array of nucleosomes is crucial to gain deeper insights into these systems.

#### Tetranucleosome array

As a final step to find suitable conformations of HUb inside chromatosomes we also applied our procedure to a tetranucleosome array published by Schalch et al. (PDB 1ZZB). [4] Similar to most chromatin complexes on the protein database, this X-ray structure omits the linker histone. We approximated theoretical positions of the 4 linker histones in this complex by protein sequence and position alignment of the core histones in the tetranucleosome array with PDB 5NL0 and the off-dyad chromatosome (Materials and Methods: Fitting HUb into the chromatosome and S5 Table). A position histogram from the atomic positions of the tetranucleosome was created to thereupon construct the scoring histogram in the same fashion as for the other nucleosomes. Subsequently, all HUb conformations were again put into each of the 4 possible linker histone positions and the corresponding ISA score was calculated Fig 7). The same regions that previously resulted in higher scores now also result in high scores for the tetranucleosome array. However, the high score region is more extended and only the conformations located in the lower left corner of the sketch-map projection should be considered as “fitting”. The “per-histone-position score distributions” (B) and (C) are created for the 4 on-dyad and off-dyad positions, respectively. The cluster with the lowest score is selected for each of the on-dyad positions and rendered in (D). The cluster centroids are marked accordingly in (A). Clusters located in positions 2 and 3 generally exhibit lower scores than 1 and 4. At this point it is not clear whether this is due to the limited length of the tetranucleosome array or whether longer nucleosome arrays exhibit more similar per-histone-position scores over their full length. To model such a system more research needs to be conducted. The packaging (on which the scores depend) of the nucleosomes is still a topic of ongoing discussion. However, our findings strongly suggest that there are native HUb conformations that fit well into the limited space available in a nucleosome array.

**Figure 7.**
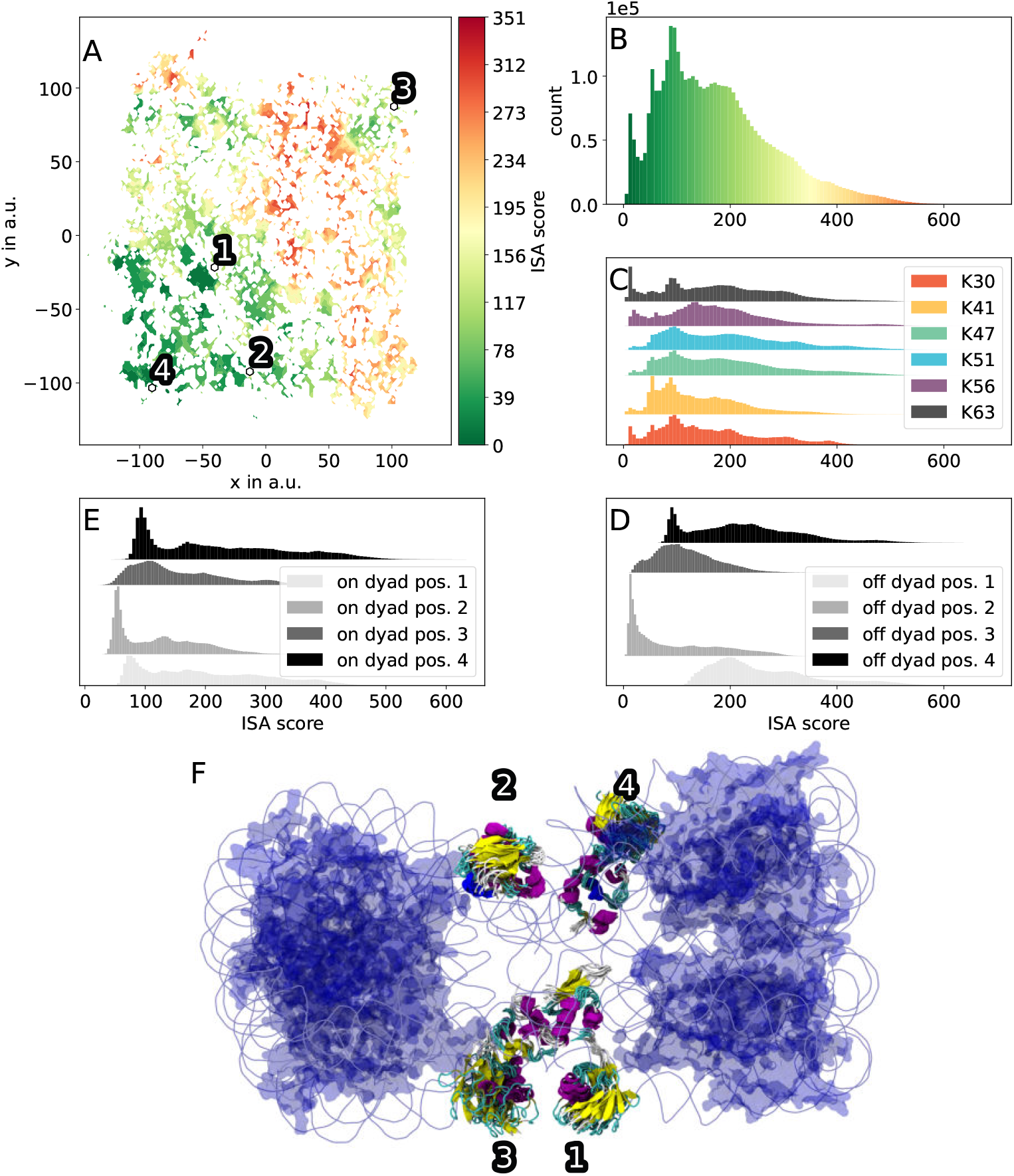
Using ISA to gauge the positions of HUb in a tetranucleosome array. The median ISA score over the 8 possible HUb positions (4 nucleosomes in the array using on-dyad and off-dyad binding motifs) exhibit large regions of higher ISA scores (A). In (B-E) the distribution of ISA scores is visualized for all 8 positions combined (B), divided by ubiquitylation site (C), and divided by the 4 off-dyad and 4 on-dyad HUb positions (E) and (D), respectively. The render in (F) uses the on-dyad 5NL0 chromatosome as reference structure for the linker histone positions 1 and 4 and the off-dyad reference structure for the positions 2 and 3. The geometric cluster centroids are annotated in (A) accordingly. They exhibit mean ISA scores between 44 and 70. These clusters are chosen for illustrative purposes to make exhibit how certain sub-states in the conformational landscape of all HUb variants can fit into the more restrictive tetranucleosome array. For the four positions 1, 2, 3, and 4, the chosen clusters contain predominantly the linkage types K51, K30, K51, and K41, respectively. The DNA is rendered as light blue tubes, core histones are blue translucent. Secondary structure elements are colored as default in VMD. [43]

Additionally to the ISA scores for the HUb variants, we looked at the position of the ubiquitin with respect to DNA. For this, all sampled conformations of the six variants were placed into PDB ID 5NL0 and the mean per-residue distance to the DNA used to color the crystal structure PDB 1UBQ (Fig 6 (I)). The α_1_ helix of the Ub-subunit exhibits shorter distances (white) to the DNA than most of the residues in the β -grasp region (blue).

## Discussion

In this work we have shown that differently ubiquitylated avian linker histones take on a wide range of conformations that are mainly dictated by the chosen ubiquitylation site although some intermixing of regions in phase space can be observed. MD simulations in combination with an expansion scheme that accelerates the exploration of phase space created a large and exhaustive library of HUb conformations. The scheme explores the relatively shallow free-energy landscape with many conformational sub-states that were identified by density based clustering on two-dimensional representation of the conformational space. These clusters represent local minima and were visited by multiple simulations from the same, sometimes even from different, topologies. The clusters pose as statistically significant (meta)stable sub-states that a real-life HUb protein might adapt. Compared to other structural methods like NMR spectroscopy or X-ray crystallography we created dynamic data of a protein modeling its time-resolved conformational changes.

We developed an interpenetration and scoring algorithm (ISA) that exploits the geometric properties of the system and shows that complex and computationally expensive scoring functions are not needed if the systems are large enough. A good correlation with pyROSETTA could be achieved by implementing cumulative 3D histograms of atomistic positions. With that we were able to identify regions of the combined HUb phase space that contain clusters that fit well into any of the 4 available chromatosome structures. Although the nucleosomes were not present during our simulations, we found roughly 93% of all conformations to be fitting into the DNA pocket of the nucleosome. This strongly indicates that the HUb protein has a natural affinity to be placed in this environment. The HUb conformations are well suited for modeling HUb inside a nucleosome and could easily be taken as input structures for further investigation via MD simulations as they generally do not need energy minimization. Such simulations don’t need to be long to probe the interactions between the Ub subunit and the DNA or the core histones.

Besides the on-dyad and off-dyad chromatosomes we applied our algorithm to a tetranucleosome array. We found that, even though the DNA-pockets of the tetranucleosome array are more restrictive, multiple clusters could easily be placed into it without significant clashes, independent whether a on-dyad or off-dyad chromatosome was used to approximate the linker histone positions.

Additionally, we found that the ubiquitin subunit in our simulations has the α_1_ -helix (beside the isopeptide bound N-terminal region) closer to the DNA than the β -grasp of Ub (Fig 7 (E)). Whether the 3 lysine residues in the α_1_ -helix could favorably interact with the DNA is certainly a worthwhile question for further examinations of this system.

Our insights into how ubiquitylated linker histone might fit into chromatosomes can help in further developing models for the stabilization of chromatin.

## Materials and Methods

### Enhanced Sampling of HUb Conformations

We created starting structures for MD simulations by linking the globular domain of avian linker histone H1 (PDB 1GHC [29]) and ubiquitin (PDB 1UBQ [44]) via an isopeptide bond between H1’s K30, K41, K47, K51, K56, and K63, residues and ubiquitin’s C-terminal glycine using the program UCSF Chimera [45] (matching lysine indices in sequence aligned human linker histone H1.2 are displayed in Fig 1). These proteins will be called K30Ub, K41Ub, K47Ub, K51Ub, K56Ub, and K63Ub, respectively. Simulations were performed for 1 to 3 different initial rotations of the lysine’s sidechain dihedral χ_3_ angle for ≈ 1 μs (for more detailed information about simulation parameters and protocol refer to S1 Text). After already ≈ 100 ns of the 1 μs simulated time the simulations of the different variants collapsed into compact distinct HUb structures, which exhibited little structural changes for the remainder of the simulations (S1 Fig). The long simulations could not decorrelate from the influence of the rotation angle χ_3_ in the starting structure even in 1 μs simulation time and became “stuck” with little variations in conformation.

To reduce bias of the starting rotation of the χ_3_ angle and push the systems to fully explore their conformational space with all-atom MD without adding biasing potentials we chose to start more rotamers of the variants by additionally varying the χ_3_ sidechain dihedral angle in 20° steps. All in all 6 × 18 = 108 starting structures were created. As the exploratory long simulations had collapsed quite rapidly, we decided on 20 ns long initial simulations from which then an expansion scheme was started that is tailored to push the system to explore new regions of conformational phase space instead of getting stuck in few collapsed structures.

This expansion scheme has already been successfully used to study the conformational space of intrinsically disordered peptides. [40] It accelerates the sampling by running many parallel simulations and assessing in regular intervals whether simulations have been trapped in low-energy basins. Identification of these basins is done by binning a low-dimensional projection of the conformational space of the HUb variants. For this reduction in dimensionality, we first needed to obtain high-dimensional descriptors of HUb that represent the protein’s conformation. For this system we chose to use the distances between the C _*α*_-atoms to provide these high-dimensional descriptors, henceforth called collective variables (CVs). A conformation of the HUb can then be represented as a collection of these distances. Instead of using the full set of 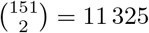 pairwise distances we chose to compare two different subsets of CVs: the solvent accessible surface area (SASA)-CVs were calculated according to Eq (1) by only using the C_*α*_ atoms with a SASA greater than 1 in the respective crystal structures. The residue-wise minimal distance collective variables (RMD-CVs) have been successfully employed to characterize di-Ubiquitin. [46] These CVs were obtained by calculating the row-wise minimum of the pairwise distance matrix between the C_*α*_ atoms of the histone and ubiquitin subunit according to Eq (2).

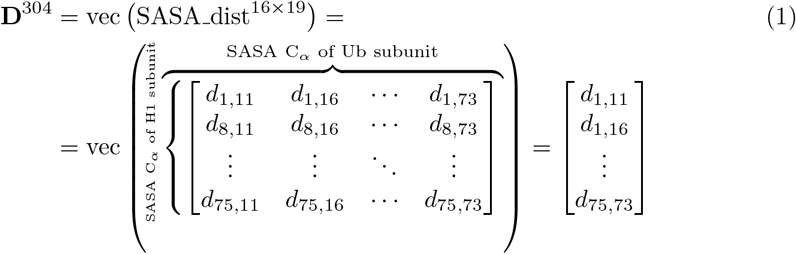

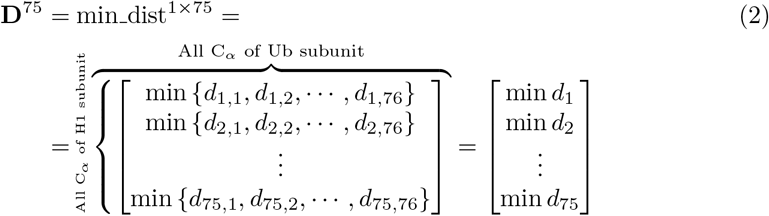

Using either of the CV datasets (K47Ub and K51Ub with SASA-CVs, K30Ub, K41Ub, K56Ub, and K63Ub with RMD-CVs) we were able to project the 18 × 20 ns simulations into a two-dimensional projection (sometimes also called map) using the sketch-map [47] method (S2 Fig, S3 Fig and S1 Text). In these projections every conformation is represented by a point; each point can be assigned to one specific structure in the conducted simulations. Note that for driving the expansion scheme, separate 2-dimensional projections of the six variants were generated (only later for analysis purposes the different systems were projected together into a combined low-dimensional map). After every 20 ns the simulations were stopped, re-projected and the expansion scheme was applied to yield 20 new starting structures. After the 3rd expansion step new landmarks were selected and the expansion scheme was continued for 6 more times (1 initial simulation run, 9 expansions) resulting in a total of ≈ 35 μs simulated time (Eq (3)).

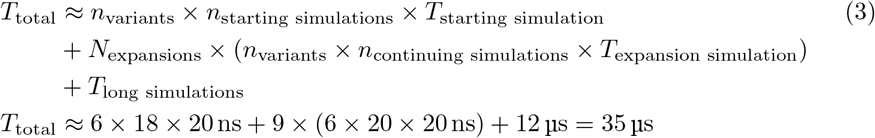

### Identifying characteristic conformational states of HUb

Subsequently all 483 983 conformations from the ≈ 35 μs aggregated simulation time were projected into a combined 2-dimensional map using the SASA-CVs to allow for analyses to work on all ubiquitination variants. A joint projection was a key argument in describing and comparing different Ub dimers by Berg et al. [46] But contrary to the projection of Ub dimers which used RMD-CVs, we found the SASA-CVs to be more suitable for a combined projection (S1 Text). From this projection, conformational states were determined with the help of a density based clustering method, HDBSCAN. [48]

HDBSCAN was first used with a minimal cluster size of 1000 to exclude projection artifacts (S1 Text). After the exclusion of these artifacts further clustering in the dimensional space was carried out iteratively to obtain structurally consistent clusters. HDBSCAN takes at least one hyperparameter for its clustering algorithm. This parameter defines the minimum cluster size to differentiate between clusters and noise points after HDBSCAN’s space transformation. We reduced the hyperparameter step-wise from large to small clusters as not every sub-state of the protein’s phase space exhibited a similar density. The minimal cluster size was set to 750, 500, 250, 125, 75, 50, and 25, respectively, in 8 clustering passes. After each pass the homogeneity of structures in the obtained clusters was checked as follows. First, a cluster’s RMSD centroid was determined by using the python package MDTraj [49]. Second, the RMSD between the centroid and the remaining cluster points was calculated. The same was done for the distance in the distance in the two-dimensional projection. Third, a set of criteria were checked:

1. A mean RMSD distance between the cluster’s RMSD centroid and the other structures greater than 6Å.
2. A Fisher-Pearson coefficient of skewness of the sketch-space distance distribution greater than 0.5.
3. Multimodality of the distribution as calculated by the python package unidip. [50]

If for any given cluster at least one of the listed conditions was satisfied, this cluster was not assigned at that clustering iteration and points belonging to it were put back into the pool of all possible points to be resolved by subsequent (smaller cluster size) clustering (an example of such a cluster is shown in Fig 8).

**Figure 8.**
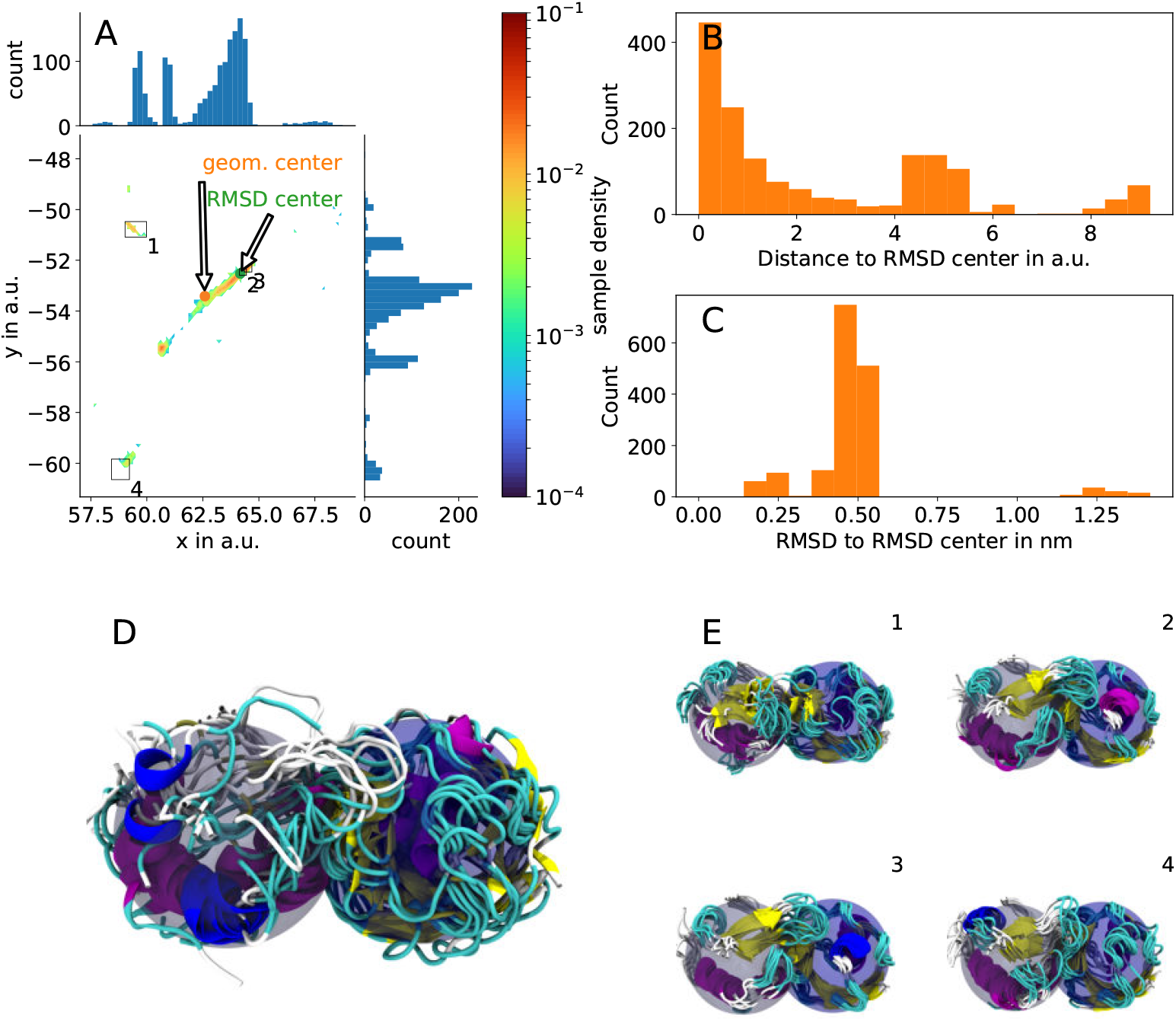
Refinement of non-uniform clusters via iterative HDBSCAN. An initial run of HDBSCAN combines all points in (A) into a single cluster. Rendering this cluster would result in a non-uniform bundle of structures (D). Furthermore, the spatial distribution in the low-dimensional space (B) and the distribution of RMSD values (C) indicate, that there are multiple sub-clusters that could be resolved by subsequent clustering. Using a lower minimal cluster size parameter the larger cluster is decomposed into smaller clusters (1-4) and the renderings of these regions (E 1-4) also show greater uniformity. Renders show the histone subunit (blue translucent sphere) on the right side and the ubiquitin subunit (grey translucent sphere) on the left side.

The resulting 710 clusters were then used as representative sub-states of HUb to be evaluated further.

### Fitting HUb into the chromatosome

To get a measure of how well a given HUb structure “fits” into a parent chromatosome we developed an algorithm using the geometric properties of the HUb chromatosome system. Our interpenetration and scoring algorithm (ISA) uses the three-dimensional histogram of atomic positions of a nucleosome as its basis. The nucleosome coordinates are taken from one of the already published chromatosome structures (Table 1). The simulated HUb structures are now positioned into the nucleosome by superimposing the histone subunit on the coordinates of the reference linker histone using RMSD minimization. The Ub subunit will now possibly intersect with some parts of the nucleosome because the nucleosome was not present during the simulations. The extent of this clash (score) is now estimated by adding-up where the Ub subunit intersects with the nucleosome. This is done by binning the atomic coordinates of the nucleosome, creating a 3D histogram of atom counts. Every atom of the Ub subunit contributes to the score with the value of the bin it intersects and the score is calculated as the sum of the per-atom bin values.

An important factor to consider for this histogram (and histograms in general) is the number of bins. The larger the number of bins, the more “empty space” will be present in the 3D histogram. This creates a problem: The smaller the size of the bins, the more zero-count bins are in the 3D histogram. However, a fine binning would be beneficial to accurately model the surface of the nucleosome. To overcome this problem, we decided to implement a cumulative histogram in the scoring process. Here, eight separate cumulative histograms starting from the eight corners of the nucleosome’s bounding box are constructed. Following the diagonal of the bounding box each bin is assigned the number of the contained atoms plus the values assigned to its preceding and neighboring bins (see S1 Text: for examples in 1D and 2D spaces). These 8 cumulative histograms are merged by assigning for every bin the respective lowest value of the 8 cumulative histograms. The resulting scoring histogram captures the surface of the nucleosome via a fine binning, while still penalizing any Ub atoms that reach into the nucleosomal core with a high score. The scoring histogram is then min-max normalized onto the range [0, 1] to make values of scores more intuitive. At the end a single value is obtained for a given HUb (ligand) and nucleosome (receptor) pose.

With this new algorithm we were able to score all 483 983 conformations sampled by MD in record time (details are discussed in Results: Scoring HUb conformations into nucleosome structures), so no pre-screening or “cherry-picking” had to be conducted. Scores were calculated by superposing a given HUb conformation using the python package MDTraj’s superpose functionality and the Cα atoms of the crystal structure’s linker histones (Table 1). [49] If the number of Cα atoms of HUb (PDB 1GHC) did not match the number of the crystal structure’s linker histone a sequence alignment with T-COFFEE [51] was carried out beforehand. Given the huge size of both the molecular system and the dataset we designed the ISA to be particularly efficient by also parallelizing it using python’s built-in joblib [52] package.

Still, a fast algorithm needs to yield usable results, so we validate our novel and purely geometric approach to scoring with tried and tested scoring functions from ROSETTA [53, 54] (we used pyROSETTA). We were able to calculate ISA scores at a speed of 140 poses per second. In contrast, pyROSETTA took ≈10 s per pose, which makes ISA roughly 1000 times faster than pyROSETTA. Although ISA does not use sophisticated potentials to model attractive and repulsive forces, a satisfying correlation of 0.88 to pyROSETTA (see Fig 9).

**Figure 9.**
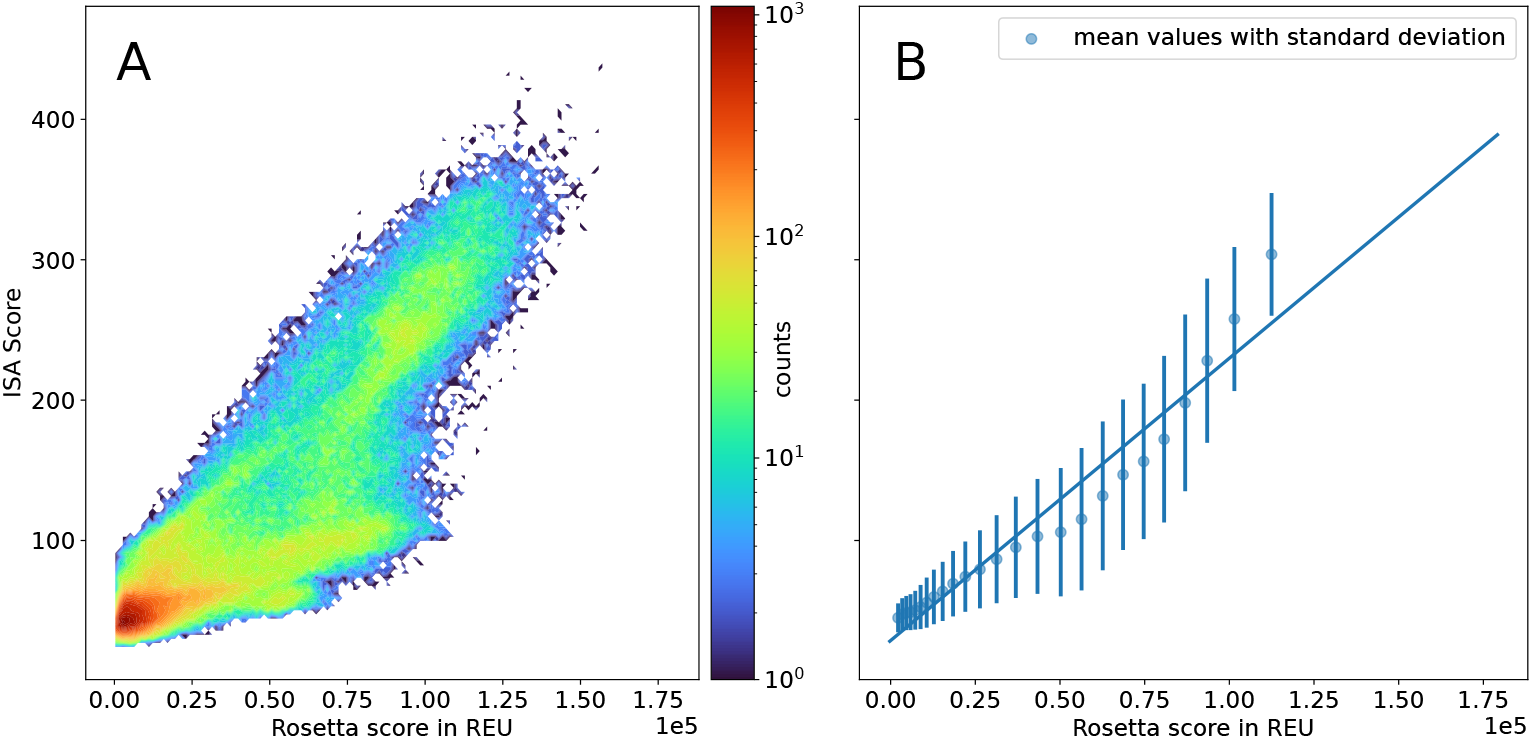
Comparison of scores obtained by our interpenetration and scoring algorithm (ISA) and ROSETTA for a subset of structures. We chose a random subset of structures from the expansion scheme simulations to score using ROSETTA and ISA. Rosetta scores range from −250 to ≈ 18 000 Rosetta Energy Units (REU). ROSETTA’s scoring functions with default weights support our claim that many structures exhibit lower scores, as the most structures can be found for lower scores where a high bin count can be observed (A). Our ISA algorithm correlates with the much more sophisticated ROSETTA algorithm with a Pearson correlation coefficient of 0.88 (B). However, our algorithm was 1000 times faster than pyROSETTA (both algorithms have been parallelized on a per-structure basis using the Python package joblib [52]).

#### Alignment of chromatosome structures into tetranucleosome

Chromatin contains many chromatosomes stacked in an orderly fashion and not just a single H1-nucleosome complex, so we also employed the ISA on a tetranucleosome array (PDB 1ZBB) published by Schalch et al. [4] This structure lacks the linker histones and thus we used a sequence and RMSD alignment of 1ZBB’s core histones with the core histones of 5NL0 [30] and the off-dyad chromatosome structure. For this we aligned the sequences of the 8 core histones with T-COFFEE and chose the longest uninterrupted residue sequence for RMSD alignment using MDTraj. Some additional adjustments had to be carried out to prevent the linker histone from clashing with the DNA of 1ZBB by manually adjusting the rotation of the chromatosomes around the third principal axis (S5 Table). Using these modelled linker histone positions we could directly use the ISA to get scores for HUb poses in the tetranucleosome array.

## Acknowledgements

Thanks to Felizitas Kirner for careful reading of the manuscript.

## Supporting information

**S1 Text. Supplementary Information**. This file contains further details about the MD simulation setup (starting structures, initial simulations, expansion scheme implementation, MD parameters, long MD trajectories), dimensionality reduction procedure with sketch-map (procedure and parameters, individual sketch-map projections and comparison of high-dimensional collective variables, high RMSD variance region), ROSETTA scoring, examples of ISA in 1D and 2D cases, overview of the 50 largest clusters, and manual adjustments to linker histone positions in 1ZBB. It contains following tables:

**S1 Table**. Long ≈ 1 μs simulations carried out to study the system.

**S2 Table**. Sketch-map parameters used to project the SASA-CVs and the RMD-CVs.

**S3 Table**. Overview of the 50 largest clusters.

**S4 Table**. Overview over the composition and scores of the ubiquitylation sites.

**S5 Table**. Manual adjustments to create hypothetical linker histone positions in the tetranucleosome array 1ZBB.

**S1 Fig. Center of geometry distances of the initial long simulations and evolution of secondary structure motifs of one of these simulations**. Time evolution of the center of geometry distances between the Ub-subunit and the H1-subunit of nine 1 μs simulations (A). Simulation names are composed of ubiquitylated lysine and starting χ_3_ angle (KXXX χ_3_). The inset figure shows the same data in a 0–100 ns interval. Raw distances are transparent. Running averages over 7.5 ns are opaque. After relaxation (until ≈ 25 ns) the center of geometry stays similar during the evolution of the whole simulation. K47Ub with χ_3_ = 20° exhibits larger variations at the beginning, but approaches an equilibrium after that. The secondary structure motifs of this simulation also exhibit convergence of the structure (B).

**S2 Fig. Sketch-map projections of the initial simulations of K30Ub and K47Ub with RMSD evolution as inset figures**. Exemplary sketch-map projections of initial simulations of K47Ub using SASA-CVs (A) and K30Ub using RMD-CVs (B). The color code indicates the starting χ_3_ angle of the respective ubiquitylated lysine residue. Similar angles have similar color saturation. Inset figures show RMSD evolution of selected trajectories. Larger RMSD deviations result in disjoint, scattered points. Smaller RMSD deviations yield cohesive patches. The simulation of K47Ub with a starting angle of χ_3_ = 0° exhibits little change in its RMSD after relaxation, which results in a densely populated patch (circle in (A)). Arrows in and annotated distances (B) connect the first and last point of the simulations with starting angle χ_3_ = 80° and χ_3_ = 120° and indicate the time evolution of these trajectories. The RMSD of the structures from K30Ub with an angle of χ_3_ = 120° exhibits a gradual increase, which is also traced in the sketch-map projection.

**S3 Fig. Individual sketch-map projections of the six variants during the expansion scheme colored by starting angle**. K47Ub and K51Ub were projected using the SASA-CVs (Eq (1)) as input CVs for sketch-map. The other 4 variants were projected using the RMD-CVs (Eq (2)). Points are colored according to starting χ_3_ angle of the ubiquitylated lysine residue. A cyclic colormap was chosen to represent the periodic nature of dihedral angles.

**S4 Fig. Comparison of RMD-CVs and SASA-CVs for the projection of the whole dataset**. For the top images (A-B) the RMD-CVs have been used as high-dimensional input for sketch-map. The lower images (C-D) were created by projecting the SASA-CVs. (A) and (C) are colored according to the ubiquitylation site of the variant. (B) and (D) are colored according to the mean RMSD distance to the subunit’s respective crystal structure (PDB ID 1GHC and 1UBQ). The RMD-CVs are not as suited as the SASA-CVs to project the conformational space of the six HUb proteins into the same low-dimensional map. More disorder for the top images can be observed than the for the bottom images, which contained the SASA-CVs in their creation. Thus the SASA-CVs were chosen to use as the high-dimensional collective variables for projecting the conformational space of all six variants into the same low-dimensional map.

**S5 Fig. Straightforward sketch-map projections of all variants using the SASA-CVs**. Besides the parameters for the sigmoid function all parameters were left at their standard value. 1000 high-dimensional landmarks were selected using sketch-map’s dimlandmark with the minmax option. Landmarks were projected into 2D using the annealing and MDS script provided in the sketch-map repository. The remaining points were projected using dimproj. The result is the projection in (A) with some points reaching out to 2 × 10^7^. We wrote a script that scales the projections and places the outliers back into the main projection area. However, using this straightforward approach, the RMSD of the structures could not be separated (B).

**S6 Fig. Extracted structures from the high-density patch in the low-dimensional projection**. In (A) the low-dimensional sketch-map projections (x and y coordinates) of all six variants using the SASA-CVs (Eq (1)) were plotted as a density map (colormap on the right). The high-density region is highlighted with a rectangle. In (B) 100 protein conformations originating from this region are shown. sketch-map was not able to separate these different conformations or even push them to the fringes of the projection map. For further analyses this region was excluded.

**S7 Fig. 1D example of cumulative histograms used in ISA**. The 1D bounding box ranges from 0 to 5. Two cumulative histograms are created from a base histogram (blue). One ascending (orange), the other one descending (green). Both are visualized as smoothed curves. The final histogram which is then used for scoring is obtained by choosing the minimal value of either of the two cumulative histograms (red).

**S8 Fig. 2D example of ISA’s cumulative histograms with more points**. In (A) 500 points have been placed in a ring with radius *r* = 3 ± *N* (*μ* = 0, *σ* = 0.2), where *N* is a standard normal distribution, around the origin. A normal histogram counting the number of points in a predefined set of bins is shown in (B), where yellow denotes highly occupied bins, purple empty bins. In (C) one of the four histograms obtained by walking from the lower left corner to the upper right is displayed. This results in some smoothing and “smearing” to the top right. In (D) the histogram for scoring is constructed by choosing the lowest value from the 4 cumulative histograms per bin. This histogram is used in ISA to determine the score of a given HUb-chromatosome pose.

**S9 Fig. Comparison of all ISA scores for monochromatosomes**. The simulations of the six linkage types (K30 to K63) were placed into the four parent chromatosomes (5NL0, 5WCU, 4QLC and the off-dyad structure). In (A) the scores of all poses is visualized. The count reaches as high as 150 000 because all ubiquitylation variants and chromatosomes are combined. The four subfigures (B-E) give the per ubiquitylation variant score for the four chromatosomes 4QLC, 5NL0, 5WCU, and off-dyad respectively. It can be seen, that K56Ub (purple) tends to display higher scores after being placed into the off-dyad chromatosome (E) than for the 4QLC chromatosome (B). The remaining six subfigures (F-K) give the per chromatosome score for the six ubiquitylation variants K30Ub, K41Ub, K47Ub, K51Ub, K56Ub, and K63Ub, respectively. Here, it can be seen that the scores for the chromatosomes 5WCU and 4QLC (green and dark purple) are very similar. They almost coincide for all ubiquitylation variants. Furthermore, K56Ub (is very unfavorably positioned for the off-dyad chromatosome (yellow).

**S10 Fig. Best fitting clusters per ubiquitylation site using 5NL0 as the parent chromatosome**. From all clusters found by iterative HDBSCAN clustering the cluster with the lowest ISA score containing at least 90% of a specific HUb variant is chosen. 20 equally spaced frames of these clusters are then placed into the parent 5NL0 chromatosome structure and visualized here. All of them have the Ub subunit pointing downward. For scores of other clusters refer to S3 Table.

## References

1. Olins AL, Olins DE. Spheroid chromatin units (ν bodies). Science. 1974;183(4122):330–332.

2. Kornberg RD. Chromatin structure: a repeating unit of histones and DNA. Science. 1974;184(4139):868–871.

3. Carruthers L, Bednar J, Woodcock C, Hansen J. Linker histones stabilize the intrinsic salt-dependent folding of nucleosomal arrays: mechanistic ramifications for higher-order chromatin folding. Biochemistry. 1998;37(42):14776—14787. doi:10.1021/bi981684e.

4. Schalch T, Duda S, Sargent DF, Richmond TJ. X-ray structure of a tetranucleosome and its implications for the chromatin fibre. Nature. 2005;436(7047):138–141.

5. Robinson PJ, Fairall L, Huynh VA, Rhodes D. EM measurements define the dimensions of the “30-nm” chromatin fiber: evidence for a compact, interdigitated structure. Proceedings of the National Academy of Sciences. 2006;103(17):6506–6511.

6. Kruithof M, Chien FT, Routh A, Logie C, Rhodes D, Van Noort J. Single-molecule force spectroscopy reveals a highly compliant helical folding for the 30-nm chromatin fiber. Nature structural & molecular biology. 2009;16(5):534–540.

7. Maeshima K, Imai R, Tamura S, Nozaki T. Chromatin as dynamic 10-nm fibers. Chromosoma. 2014;123(3):225–237.

8. Norouzi D, Zhurkin VB. Topological polymorphism of the two-start chromatin fiber. Biophysical journal. 2015;108(10):2591–2600.

9. Luger K, Dechassa ML, Tremethick DJ. New insights into nucleosome and chromatin structure: an ordered state or a disordered affair? Nature reviews Molecular cell biology. 2012;13(7):436–447.

10. Zhou BR, Feng H, Kato H, Dai L, Yang Y, Zhou Y, et al. Structural insights into the histone H1-nucleosome complex. Proceedings of the National Academy of Sciences. 2013;110(48):19390–19395.

11. Bednar J, Hamiche A, Dimitrov S. H1–nucleosome interactions and their functional implications. Biochimica et Biophysica Acta (BBA) - Gene Regulatory Mechanisms. 2016;1859(3):436 – 443. doi:https://doi.org/10.1016/j.bbagrm.2015.10.012.

12. Sun ZW, Allis CD. Ubiquitination of histone H2B regulates H3 methylation and gene silencing in yeast. Nature. 2002;418(6893):104–108.

13. Zentner GE, Henikoff S. Regulation of nucleosome dynamics by histone modifications. Nature structural & molecular biology. 2013;20(3):259.

14. Kim W, Bennett EJ, Huttlin EL, Guo A, Li J, Possemato A, et al. Systematic and quantitative assessment of the ubiquitin-modified proteome. Molecular cell. 2011;44(2):325–340.

15. Wagner SA, Beli P, Weinert BT, Nielsen ML, Cox J, Mann M, et al. A proteome-wide, quantitative survey of in vivo ubiquitylation sites reveals widespread regulatory roles. Molecular & Cellular Proteomics. 2011;10(10).

16. Danielsen JM, Sylvestersen KB, Bekker-Jensen S, Szklarczyk D, Poulsen JW, Horn H, et al. Mass spectrometric analysis of lysine ubiquitylation reveals promiscuity at site level. Molecular & Cellular Proteomics. 2011;10(3).

17. Chang L, Shen L, Zhou H, Gao J, Pan H, Zheng L, et al. ITCH nuclear translocation and H1. 2 polyubiquitination negatively regulate the DNA damage response. Nucleic acids research. 2019;47(2):824–842.

18. Thorslund T, Ripplinger A, Hoffmann S, Wild T, Uckelmann M, Villumsen B, et al. Histone H1 couples initiation and amplification of ubiquitin signalling after DNA damage. Nature. 2015;527(7578):389–393.

19. Ding X, Lin X, Zhang B. Stability and folding pathways of tetra-nucleosome from six-dimensional free energy surface. Nature communications. 2021;12(1):1–9.

20. Sharma S, Ding F, Dokholyan NV. Multiscale modeling of nucleosome dynamics. Biophysical journal. 2007;92(5):1457–1470.

21. Arya G, Schlick T. Role of histone tails in chromatin folding revealed by a mesoscopic oligonucleosome model. Proceedings of the National Academy of Sciences. 2006;103(44):16236–16241.

22. Brandani GB, Tan C, Takada S. The kinetic landscape of nucleosome assembly: a coarse-grained molecular dynamics study. PLoS computational biology. 2021;17(7):e1009253.

23. Huertas J, Schöler HR, Cojocaru V. Histone tails cooperate to control the breathing of genomic nucleosomes. PLoS computational biology. 2021;17(6):e1009013.

24. Allan J, Hartman P, Crane-Robinson C, Aviles F. The structure of histone H1 and its location in chromatin. Nature. 1980;288(5792):675–679.

25. Pruss D, Bartholomew B, Persinger J, Hayes J, Arents G, Moudrianakis EN, et al. An asymmetric model for the nucleosome: a binding site for linker histones inside the DNA gyres. Science. 1996;274(5287):614–617.

26. Zhou YB, Gerchman SE, Ramakrishnan V, Travers A, Muyldermans S. Position and orientation of the globular domain of linker histone H5 on the nucleosome. Nature. 1998;395(6700):402–405.

27. Syed SH, Goutte-Gattat D, Becker N, Meyer S, Shukla MS, Hayes JJ, et al. Single-base resolution mapping of H1–nucleosome interactions and 3D organization of the nucleosome. Proceedings of the National Academy of Sciences. 2010;107(21):9620–9625.

28. Woods DC, Wereszczynski J. Elucidating the influence of linker histone variants on chromatosome dynamics and energetics. Nucleic acids research. 2020;48(7):3591–3604.

29. Cerf C, Lippens G, Muyldermans S, Segers A, Ramakrishnan V, Wodak SJ, et al. Homo-and heteronuclear two-dimensional NMR studies of the globular domain of histone H1: sequential assignment and secondary structure. Biochemistry. 1993;32(42):11345–11351.

30. Bednar J, Garcia-Saez I, Boopathi R, Cutter AR, Papai G, Reymer A, et al. Structure and dynamics of a 197 bp nucleosome in complex with linker histone H1. Molecular cell. 2017;66(3):384–397.

31. Höllmüller E, Geigges S, Niedermeier ML, Kammer KM, Kienle SM, Rösner D, et al. Site-specific ubiquitylation acts as a regulator of linker histone H1. Nature Communications. 2021;12(1):1–15.

32. Fierz B, Chatterjee C, McGinty RK, Bar-Dagan M, Raleigh DP, Muir TW. Histone H2B ubiquitylation disrupts local and higher-order chromatin compaction. Nature chemical biology. 2011;7(2):113–119.

33. Höllmüller E, Greiner K, Kienle SM, Scheffner M, Marx A, Stengel F. Interactome of Site-Specifically Acetylated Linker Histone H1. Journal of Proteome Research. 2021;20(9):4443–4451.

34. Thåström A, Bingham L, Widom J. Nucleosomal locations of dominant DNA sequence motifs for histone–DNA interactions and nucleosome positioning. Journal of molecular biology. 2004;338(4):695–709.

35. Correll SJ, Schubert MH, Grigoryev SA. Short nucleosome repeats impose rotational modulations on chromatin fibre folding. The EMBO journal. 2012;31(10):2416–2426.

36. Rösner D. Chemical mono-ubiquitylation of linker histone H1.2 by combining unnatural amino acids with click chemistry: Synthesis and structural studies [Ph.D. thesis]. University of Konstanz. Universitä tsstr. 10, D-78464 Konstanz, Germany; 2015. Available from: http://nbn-resolving.de/urn:nbn:de:bsz:352-0-326144.

37. Ramakrishnan V, Finch J, Graziano V, Lee P, Sweet R. Crystal structure of globular domain of histone H5 and its implications for nucleosome binding. Nature. 1993;362(6417):219–223.

38. Zhou BR, Jiang J, Feng H, Ghirlando R, Xiao TS, Bai Y. Structural mechanisms of nucleosome recognition by linker histones. Molecular cell. 2015;59(4):628–638.

39. Zhou BR, Jiang J, Ghirlando R, Norouzi D, Yadav KS, Feng H, et al. Revisit of reconstituted 30-nm nucleosome arrays reveals an ensemble of dynamic structures. Journal of molecular biology. 2018;430(18):3093–3110.

40. Kukharenko O, Sawade K, Steuer J, Peter C. Using dimensionality reduction to systematically expand conformational sampling of intrinsically disordered peptides. Journal of Chemical Theory and Computation. 2016;12(10):4726–4734.

41. Berg A, Peter C. Simulating and analysing configurational landscapes of protein–protein contact formation. Interface focus. 2019;9(3):20180062.

42. Arya G, Zhang Q, Schlick T. Flexible histone tails in a new mesoscopic oligonucleosome model. Biophysical journal. 2006;91(1):133–150.

43. Humphrey W, Dalke A, Schulten K. VMD: visual molecular dynamics. Journal of molecular graphics. 1996;14(1):33–38.

44. Vijay-Kumar S, Bugg CE, Cook WJ. Structure of ubiquitin refined at 1.8Åresolution. Journal of molecular biology. 1987;194(3):531–544.

45. Pettersen EF, Goddard TD, Huang CC, Couch GS, Greenblatt DM, Meng EC, et al. UCSF Chimera—a visualization system for exploratory research and analysis. Journal of computational chemistry. 2004;25(13):1605–1612.

46. Berg A, Kukharenko O, Scheffner M, Peter C. Towards a molecular basis of ubiquitin signaling: A dual-scale simulation study of ubiquitin dimers. PLOS Computational Biology. 2018;14(11):1–14. doi:10.1371/journal.pcbi.1006589.

47. Ceriotti M, Tribello GA, Parrinello M. Simplifying the representation of complex free-energy landscapes using sketch-map. Proceedings of the National Academy of Sciences. 2011;108(32):13023–13028.

48. McInnes L, Healy J, Astels S. hdbscan: Hierarchical density based clustering. The Journal of Open Source Software. 2017;2(11). doi:10.21105/joss.00205.

49. McGibbon RT, Beauchamp KA, Harrigan MP, Klein C, Swails JM, Hernández CX, et al. MDTraj: A Modern Open Library for the Analysis of Molecular Dynamics Trajectories. Biophysical Journal. 2015;109(8):1528 – 1532. doi:10.1016/j.bpj.2015.08.015.

50. Maurus S, Plant C. Skinny-dip: clustering in a sea of noise. In: Proceedings of the 22nd ACM SIGKDD international conference on Knowledge discovery and data mining; 2016. p. 1055–1064.

51. Notredame C, Higgins DG, Heringa J. T-Coffee: A novel method for fast and accurate multiple sequence alignment. Journal of molecular biology. 2000;302(1):205–217.

52. Joblib Development Team. Joblib: running Python functions as pipeline jobs; 2020. Available from: https://joblib.readthedocs.io/.

53. Leaver-Fay A, Tyka M, Lewis SM, Lange OF, Thompson J, Jacak R, et al. ROSETTA3: an object-oriented software suite for the simulation and design of macromolecules. In: Methods in enzymology. vol. 487. Elsevier; 2011. p. 545–574.

54. Kaufmann KW, Lemmon GH, DeLuca SL, Sheehan JH, Meiler J. Practically useful: what the Rosetta protein modeling suite can do for you. Biochemistry. 2010;49(14):2987–2998.

