## Supplementary text, tables, figures for "Combining molecular dynamics simulations and scoring method to computationally model ubiquitylated linker histones in chromatosomes"

#### S1 MD simulation set up

##### S1.1 Starting structures

###### S1.1.1 Initial long simulations

Starting structures for all simulations were generated by joining ubiquitin's C-terminus (G76 of PDB ID 1UBQ [1]) with K30, K41, K47, K51, K56, and K60 of avian linker histone H1 (PDB ID 1GHC [2]) using the program UCSF Chimera. [3] The connecting lysine and glycine were renamed to LYQ and GLQ, respectively. Two of lysine's hydrogens and one oxygen atom of glycine were deleted. For the long ≈ 1 µs simulations the $\chi_3$-angle of the ubiquitylated lysine was adjusted to the values given in Table S1.

**Table S1.** Long ≈ 1 µs simulations carried out to study the system. After these simulations it became apparent, that the expansion scheme would be applicable here to generate more conformations in a shorter simulation time.

| Ubiquitylation Site | $\chi_3$ rotation in degree | Time in ps |
|---|---|---|
| K47 | -100 | 1000000 |
| K47 | 140 | 1000000 |
| K47 | 20 | 1000000 |
| K51 | -120 | 1000000 |
| K51 | 0 | 1000000 |
| K51 | 120 | 1000000 |
| K56 | -120 | 1000000 |
| K56 | 0 | 1000000 |
| K56 | 120 | 1000000 |
| K63 | -100 | 1000000 |
| K63 | 140 | 1000000 |
| K63 | 20 | 1076839 |
| K63 | 60 | 1076839 |

##### S1.1.2 Starting structures for expansion scheme

The $\chi_3$-angle was adjusted in 20° steps to create 18 starting structures for each of the ubiquitylation sites K30, K41, K47, K51, K56, and K63. Some dihedral angles resulted in starting structures that could not be energy-minimized due to clashes of the two subunits. These starting angles were omitted, which led to some specific ubiquitylation-site angle combinations not being present in the simulation set. These initial simulations then were fed into the expansion scheme. For every variant 20 new simulations from points selected in sketch-map space were started. These expansion simulations were started from the solvated system and underwent a short (0.2 ns) position-restrained simulation before conducting MD simulations, using the same parameters as in Section S1.2.

#### S1.2 MD Parameters

The starting structures were fed into all-atom molecular dynamics simulations using the GROMACS program package. [4] The GROMOS54a7 [5] forcefield was modified to include the isopeptide connection (see Section S1.2.1). The starting structures were put into a dodecahederal box with 0.8 nm distance to the walls, reducing the number of explicit solvent molecules needing to be simulated. Periodic boundary conditions were implemented, proteins were solvated in SPC water [6], and charges were neutralized with $Na^+$ and $Cl^-$ ions. The systems underwent standard steepest descent energy minimization and were equilibrated with the positions of the heavy atoms constrained. MD simulations employed the leapfrog integrator [7] with an integration step of 2 fs. The temperature was kept at 300 K using the velocity rescale algorithm [8] with a time constant of $\tau_t = 1.0$ ps. Pressure was kept at 101 325 Pa with the Berendsen thermostat [9] with a time constant of $\tau_p = 1.0$ ps. Bond constraints were handled by the LINCS algorithm. [10] Long range electrostatics were calculated by the Particle Mesh Ewald Method [11] with a Fourier spacing of 0.12 nm and cubic interpolation. Coulomb, Van-der-Waals and the long neighbor list cutoffs were set to be 1.4 nm. The short neighbor list cutoff was set to 1 nm. Neighbor lists were updated every 10 steps. The simulations were carried out on the HPC resources bwUniCluster and bwForCluster MLS&WISO.Completed simulations were stripped of solvent and frames were extracted from the trajectory every 50 ps resulting in 401 frames per trajectory.

##### S1.2.1 Adding isopeptide bonds to the GROMOS54a7 forcefield

The GROMOS54a7 forcefield was changed following a procedure described by Berg et al. [12] The renamed lysine (LYQ) and glycine (GLQ) were added to the forcefield's `residuetypes.dat` and to `aminoacids.hdb`. Additionally, the sidechain nitrogen was renamed to NQ, because it needed different interaction potentials, than lysine "standard" nitrogen NZ. The isopeptide bond between LYQ-NQ and GLQ-C was defined in the forcefield's `specbond.dat`. The two new aminoacids were added to `aminoacids.rtp` using their non-iospeptide counterparts as base and removing unwanted atoms bonds, angles, dihedrals, and impropers. The termini of COQ were adjusted in `aminoacids.c.tdb`. The proteins were built with UCSF Chimera [3] and saved as pdb files. GROMACS' `pdb2gmx` program was used with the modified forcefield to create GROMACS topologies.

#### S1.3 Slow conformational exploration of initial long simulations

The $\approx 1\,\mu s$ simulations described in Table S1 did not result in a sufficiently high sampling rate of the conformational space available to the variants (Fig S1). For this reason we decided to start more simulations and apply the expansion scheme to this system. The simulations obtained by this method exhibit not only greater variance in the distance between the center of geometry of the two subunits, but also greater distances which indicates that rare events occur more frequently.

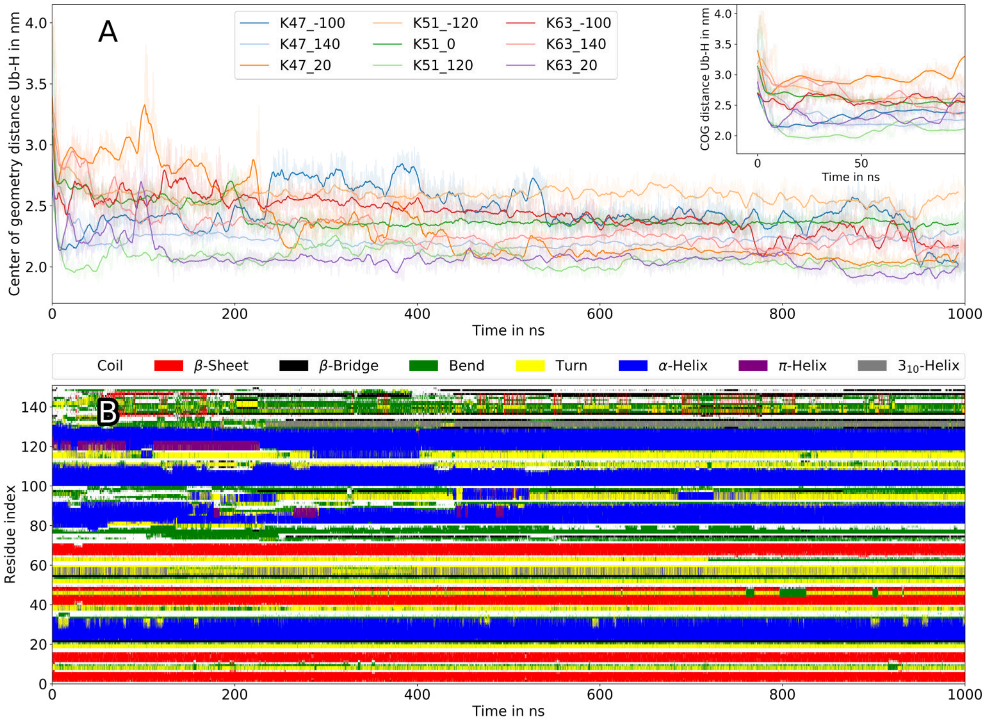


**Figure S1.** Time evolution of the center of geometry distances between the Ub-subunit and the H1-subunit of nine $1\,\mu s$ simulations (A). Simulation names are composed of ubiquitylated lysine and starting $\chi_3$ angle (KXXX_$\chi_3$). The inset figure shows the same data in a 0–100 ns interval. Raw distances are transparent. Running averages over 7.5 ns are opaque. After relaxation (until $\approx 25$ ns) the center of geometry stays similar during the evolution of the whole simulation. K47Ub with $\chi_3 = 20°$ exhibits larger variations at the beginning, but approaches an equilibrium after that. The secondary structure motifs of this simulation also exhibit convergence of the structure (B).

#### S2 Dimensionality reduction with sketch-map

##### S2.1 General procedure

We explored two different CVs as input for dimensionality reduction with sketch-map [13]. K47Ub and K51Ub were plugged into the expansion scheme using the SASA-CVs, while from K30Ub, K41Ub, K56Ub, and K63Ub the RMD-CVs were used as high-dimensional input CVs for sketch-map. The parameters of the high- and low-dimensional sigmoid functions used to transform the high- and low-dimensional CV-space and sketch-map space are given in Table S2

**Table S2.** Sketch-map parameters used to project the SASA-CVs and the RMD-CVs.

| High-D data | $D$ | $d$ | $\sigma$ | $A$ | $B$ | $a$ | $b$ |
|---|---|---|---|---|---|---|---|
| SASA-CVs | 304 | 2 | 6.0 | 10 | 3 | 2 | 3 |
| RMD-CVs | 75 | 2 | 1.5 | 5 | 2 | 1 | 2 |

For projecting the simulations of each variant separately, 1000 landmarks were selected using a farthest point sampling (`dimlandmark`). For the combined projection of all variants, 2000 landmarks were selected from the complete SASA-CV feature space. These landmarks were projected into 2D using sketch-map's simulated annealing script. One should note, that this iterative optimization procedure, is not always able to produce an optimal (or satisfactory) output for such high-dimensional and diverse data set. We have encountered two types of flaws. First, some points (from a total of 483 983 points 1306 were placed beyond the 150 a.u. of the main projection area, 357 were placed beyond 1000 a.u. and 5 were placed beyond 1 000 000 a.u.) were placed outside of the main projection area. For each of these points we selected the closest neighbor in the main projection area and placed near it at a random position within ±0.5 length units. Second, we also encountered a considerable number of points with identical (x, y) coordinates, which formed quite sharp density peaks. The structures inside those areas were very similar, but for the consistency of the analysis (without removing them density-based clustering is not able to find clusters), we removed those points from the data set for clustering and then reassigned the resulting cluster memberships.

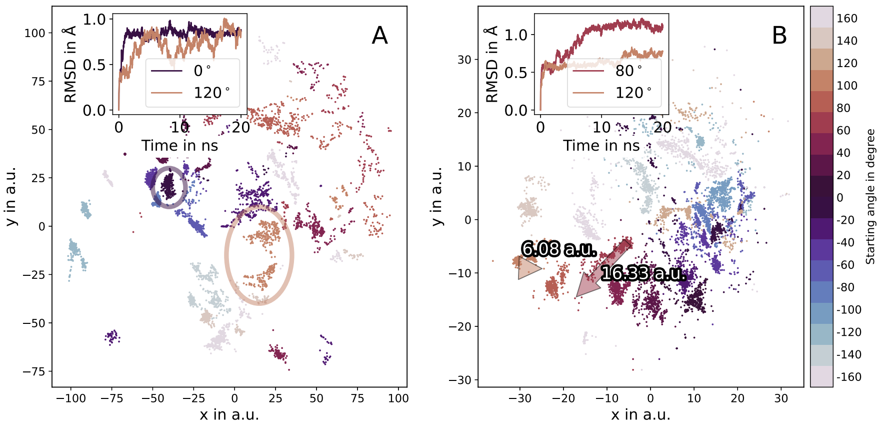


**Figure S2.** Exemplary sketch-map projections of initial simulations of K47Ub using SASA-CVs (A) and K30Ub using RMD-CVs (B). The color code indicates the starting $\chi_3$ angle of the respective ubiquitylated lysine residue. Similar angles have similar color saturation. Inset figures show RMSD evolution of selected trajectories. Larger RMSD deviations result in disjoint, scattered points. Smaller RMSD deviations yield cohesive patches. The simulation of K47Ub with a starting angle of $\chi_3 = 0°$ exhibits little change in its RMSD after relaxation, which results in a densely populated patch (circle in (A)). Arrows in and annotated distances (B) connect the first and last point of the simulations with starting angle $\chi_3 = 80°$ and $\chi_3 = 120°$ and indicate the time evolution of these trajectories. The RMSD of the structures from K30Ub with an angle of $\chi_3 = 120°$ exhibits a gradual increase, which is also traced in the sketch-map projection.

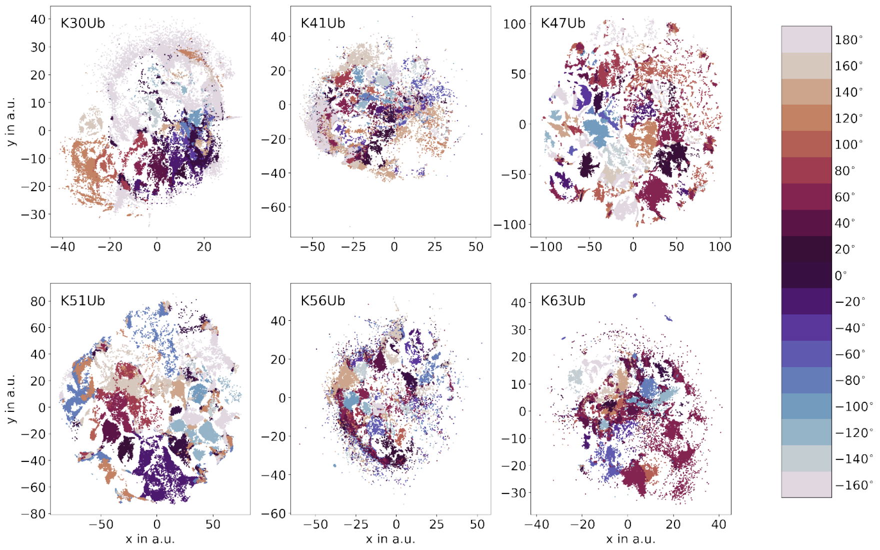


**Figure S3.** Scatter plots of individual sketch-map projections of the six variants during the expansion scheme. K47Ub and K51Ub were projected using the SASA-CVs as input CVs for sketch-map. The other 4 variants were projected using the RMD-CVs. Points are colored according to starting $\chi_3$ angle of the ubiquitylated lysine residue. A cyclic colormap was chosen to represent the periodic nature of dihedral angles.

#### S2.2 Individual sketch-map projections and comparison of high-dimensional collective variables

From the accumulated simulations a combined sketch-map projection was created using the SASA-CVs as high-dimensional input. The reason we decided against the RMD-CVs is visualized in Fig S4. Using the RMD CVs, sketch-map was not able to separate the structures in the low-dimensional projection. This is especially drastic when comparing the RMSD distance to the crystal structure (Fig S4 (B) and (C)).

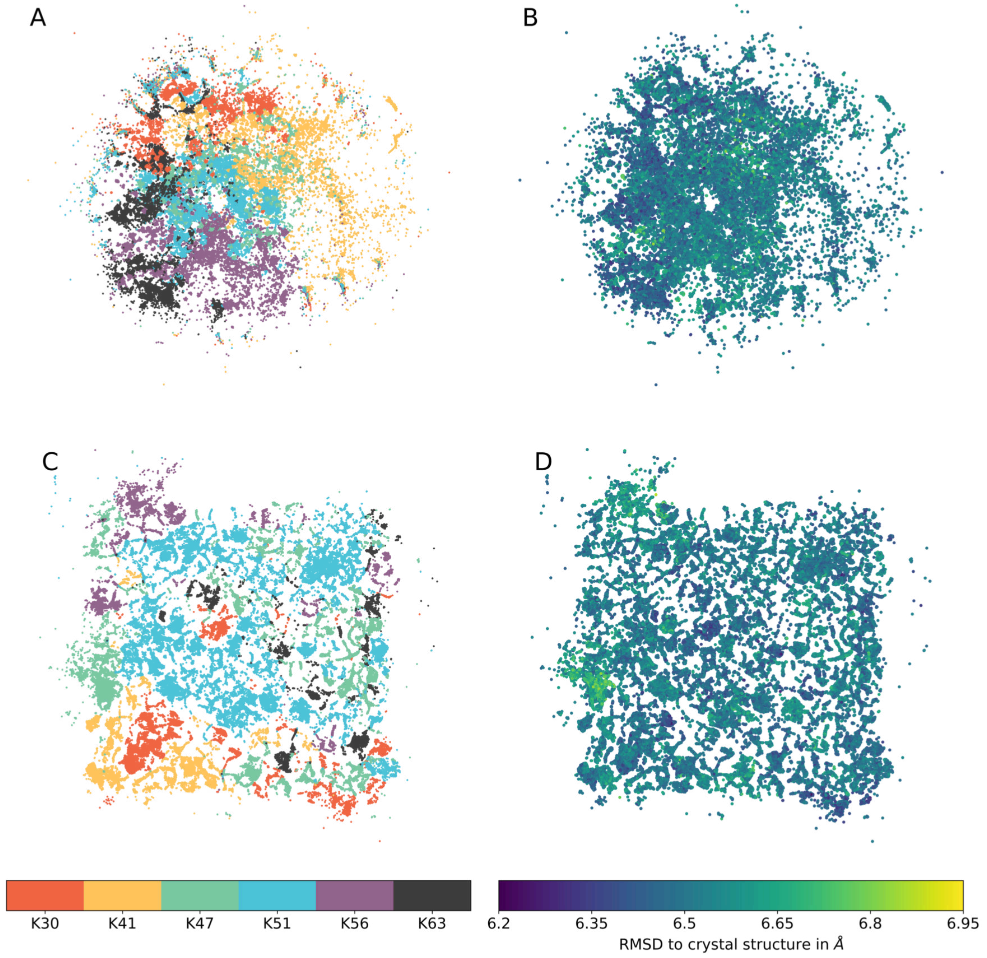


**Figure S4.** Comparison of RMD-CVs and SASA-CVs for the projection of the whole dataset. For the top images (A-B) the RMD-CVs have been used as high-dimensional input for sketch-map. The lower images (C-D) were created by projecting the SASA-CVs. (A) and (C) are colored according to the ubiquitylation site of the variant. (B) and (D) are colored according to the mean RMSD distance to the subunit's respective crystal structure (PDB ID 1GHC and 1UBQ). The RMD-CVs are not as suited as the SASA-CVs to project the conformational space of the six HUb proteins into the same low-dimensional map. More disorder for the top images can be observed than the for the bottom images, which contained the SASA-CVs in their creation. Thus the SASA-CVs were chosen to use as the high-dimensional collective variables for projecting the conformational space of all six variants into the same low-dimensional map.

#### S2.3 High RMSD variance region

Although the SASA-CVs were more suited for the projection of the combined phase space than the RMD-CVs, we could still identify some caveats: the projection of all sampled conformations into a combined sketch-map projection (similar to the individual projections (Section S2.1) was highly dependent on the chosen landmarks and oftentimes landmarks did not yield satisfactory results at all (Fig S5).

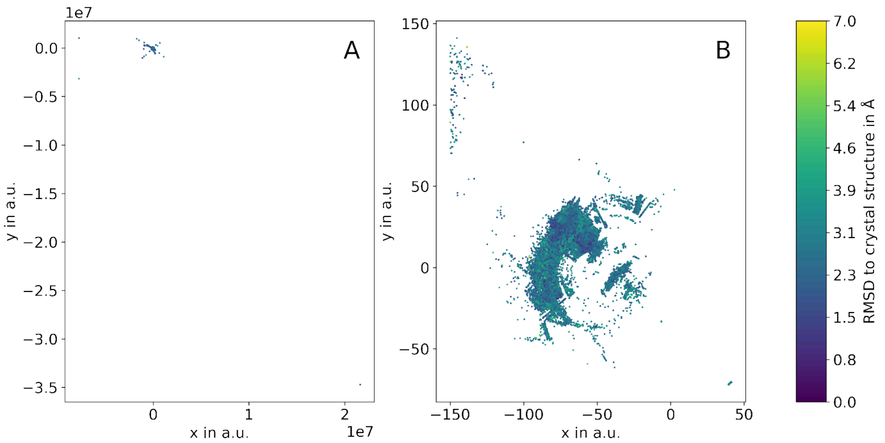


**Figure S5.** Straightforward sketch-map projections of all variants using the SASA-CVs. Besides the parameters for the sigmoid function all parameters were left at their standard value. 1000 high-dimensional landmarks were selected using sketch-map's `dimlandmark` with the minmax option. Landmarks were projected into 2D using the annealing and MDS script provided in the sketch-map repository. The remaining points were projected using `dimproj`. The result is the projection in (A) with some points reaching out to $2 \times 10^7$. We wrote a script that scales the projections and places the outliers back into the main projection area. However, using this straightforward approach, the RMSD of the structures could not be separated (B).

Furthermore, due to the amount of different structures and high-dimensionality of the initial space the sketch-map optimization was not always able to move the unique structures to the fringe regions of the landscape like it did when every variant is projected into its own low-dimensional landscape (Fig S3). In our most promising projection these unique structure can be found aggregated in a region of high RMSD variance (Section S2.3). Due to optimisation problems the sketch-map was placing some points at the exact same xy-coordinates. With this dataset, no density-based clustering could be conducted, as these xy-coordinate-duplicates would have been regarded as the density maximum and all other points would have been assigned as noise. Example of such high-density patch is shown in Fig S6. Thus, the data set made it necessary to first exclude the region of high density and high RMSD variance, then the xy-coordinate duplicates, run the density-based clustering before reassigning the xy-coordinate duplicates and assigning the high density, high RMSD variance region to the noise.

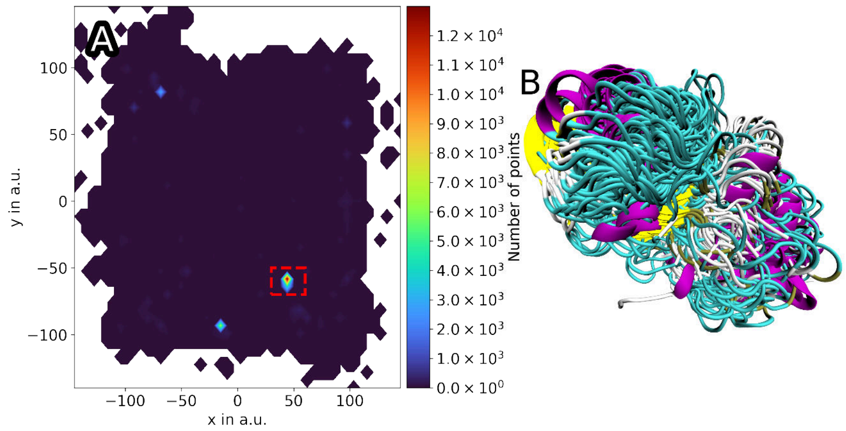


**Figure S6.** Extracted structures from the high-density patch in the low-dimensional projection. In (A) the low-dimensional sketch-map projections (x and y coordinates) of all six variants using the SASA-CVs were plotted as a density map (colormap on the right). The high-density region is highlighted with a rectangle. In (B) 100 protein conformations originating from this region are shown. sketch-map was not able to separate these different conformations or even push them to the fringes of the projection map. For further analyses this region was excluded.

### S3 ROSETTA scoring

For ROSETTA two input structures needed to be created. The receptor.pdb was created by removing the linker histone from the chromatosomes listed in the results section of the main text. The ligand.pdb was cleaned from the isopeptide LYQ and GLQ residues, because they would have caused errors in ROSETTA, and superposed using MDTraj. A scoring protocol (no dynamics, no refinement) was adapted from Evan H. Baugh's D020_Pose_scoring tutorial. [14] The weights of ROSETTA's scoring functions were left at their standard values. The comparison of ISA and ROSETTA was done on the 5NL0 chromatosome.

### S4 Examples of ISA in 1D and 2D cases

To get a better idea how the cumulative histograms in ISA are calculated and which values are finally used in scoring we will visualize the procedure in 1D and 2D here.

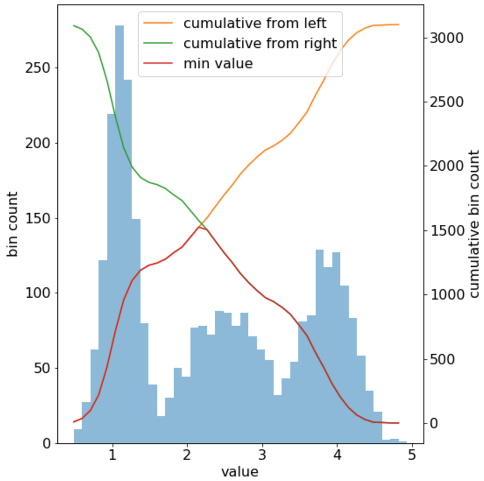


**Figure S7.** 1D example of cumulative histograms used in ISA. The 1D bounding box ranges from 0 to 5. Two cumulative histograms are created from a base histogram (blue). One ascending (orange), the other one descending (green). Both are visualized as smoothed curves. The final histogram which is then used for scoring is obtained by choosing the minimal value of either of the two cumulative histograms (red).

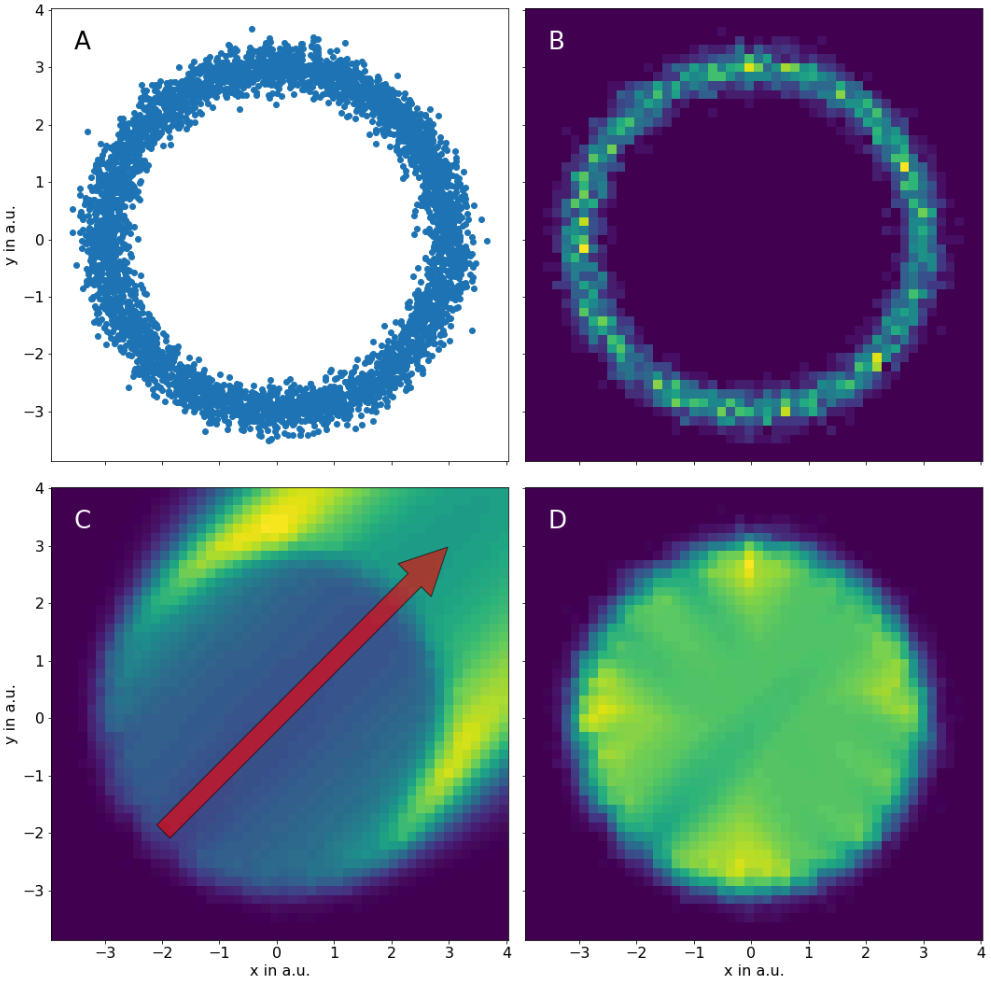


**Figure S8.** 2D example of ISA's cumulative histograms with more points. In (A) 500 points have been placed in a ring with radius $r = 3 \pm N(\mu = 0, \sigma = 0.2)$, where $N$ is a standard normal distribution, around the origin. A normal histogram counting the number of points in a predefined set of bins is shown in (B), where yellow denotes highly occupied bins, purple empty bins. In (C) one of the four histograms obtained by walking from the lower left corner to the upper right is displayed. This results in some smoothing and "smearing" to the top right. In (D) the histogram for scoring is constructed by choosing the lowest value from the 4 cumulative histograms per bin. This histogram is used in ISA to determine the score of a given HUb-chromatosome pose.

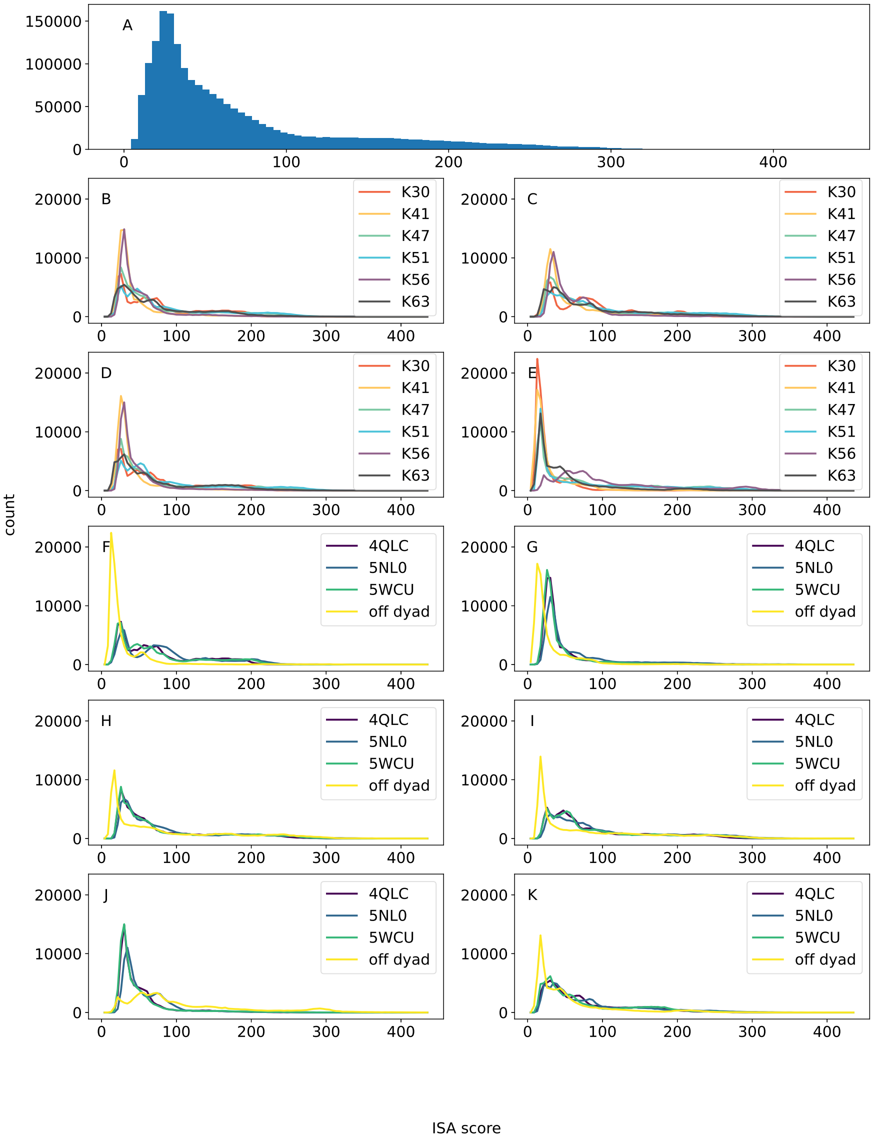


**Figure S9.** Comparison of all ISA scores for monochromatosomes. The simulations of the six linkage types (K30 to K63) were placed into the four parent chromatosomes (5NL0, 5WCU, 4QLC and the off-dyad structure). In (A) the scores of all poses is visualized. The count reaches as high as 150 000 because all ubiquitylation variants and chromatosomes are combined. The four subfigures (B-E) give the per ubiquitylation variant score for the four chromatosomes 4QLC, 5NL0, 5WCU, and off-dyad respectively. It can be seen, that K56Ub (purple) tends to display higher scores after placed into the off-dyad chromatosome (E) than for the 4QLC chromatosome (B). The remaining six subfigures (F-K) give the per chromatosome score for the six ubiquitylation variants K30Ub, K41Ub, K47Ub, K51Ub, K56Ub, and K63Ub, respectively. Here, it can be seen that the scores for the chromatosomes 5WCU and 4QLC (green and dark purple) are very similar. They almost coincide for all ubiquitylation variants. Furthermore, K56Ub (is very unfavorably positioned for the off-dyad chromatosome (yellow).

### S5 Overview of the 50 largest clusters

**Table S3.** Overview of the 50 largest clusters. The *hdbscan id* refers to the cluster id, assigned by the HDBSCAN algorithm. *cluster size* refers to the number of actual HUb conformations in this cluster. The percentages for the different linkage types give the composition of this cluster dependent on the HUb variants. *% of combined ensemble* relates the number of HUb conformations in this cluster with all simulated/sampled conformations. Following are values for ISA and ROSETTA scores. The internal RMSD was calculated by first finding the RMSD centroid of a cluster by finding the argmin of a pairwise RMSD distance matrix and then choosing that centroid as a reference for the RMSD calculations of internal RMSD. Given here are the mean (µ) and standard deviation (σ) in nm. The last column *Ub direction* gives the direction of the Ub subunit after placing HUb into an on-dyad chromatosome. For example in **??** (D), (F), and (G), the Ub subunit is pointing downwards.

| cluster num by count | hdbscan id | cluster size | % K30 | % K41 | % K47 | % K51 | % K56 | % K63 | % of combined ensemble | µ ISA score | µ ROSETTA score in REU | σ ISA score | σ ROSETTA score in REU | µ internal rmsd in nm | σ internal rmsd in nm | Ub direction |
|---|---|---|---|---|---|---|---|---|---|---|---|---|---|---|---|---|
| 1 | 11 | 3272 | 100.0 | | | | | | 0.68 | 24.95 | 5638 | 21.79 | 2026 | 0.26 | 0.04 | ↓ |
| 2 | 52 | 2691 | 0.6 | | | | | 99.4 | 0.56 | 85.23 | 72823 | 59.29 | 10468 | 0.37 | 0.10 | ↑ |
| 3 | 5 | 2417 | 100.0 | | | | | | 0.50 | 26.17 | 2777 | 35.49 | 1790 | 0.42 | 0.15 | ↓ |
| 4 | 6 | 2168 | | 100.0 | | | | | 0.45 | 51.81 | 60951 | 51.49 | 25128 | 0.56 | 0.36 | ↓ |
| 5 | 1 | 1756 | 100.0 | | | | | | 0.36 | 22.56 | 4083 | 6.50 | 2419 | 0.26 | 0.05 | ↓ |
| 6 | 45 | 1568 | | 100.0 | | | | | 0.32 | 30.28 | 18232 | 15.93 | 11640 | 0.45 | 0.23 | ↓ |
| 7 | 32 | 1466 | | | | 100.0 | | | 0.30 | 158.32 | 110874 | 90.34 | 9241 | 0.36 | 0.09 | ↑ |
| 8 | 0 | 1434 | | | | 100.0 | | | 0.30 | 58.29 | 45310 | 21.73 | 10089 | 0.28 | 0.05 | ↓ |
| 9 | 12 | 1383 | 0.1 | | | 0.4 | | 99.4 | 0.29 | 117.28 | 73411 | 47.11 | 6278 | 0.32 | 0.09 | ↓ |
| 10 | 49 | 1287 | | | | 100.0 | | | 0.27 | 107.12 | 75174 | 70.28 | 19051 | 0.30 | 0.08 | ↓ |
| 11 | 25 | 1103 | | | | 100.0 | | | 0.23 | 159.23 | 99367 | 99.85 | 10012 | 0.28 | 0.05 | ↑ |
| 12 | 158 | 1094 | | | 100.0 | | | | 0.23 | 22.02 | 6673 | 14.35 | 2823 | 0.33 | 0.06 | ↓ |
| 13 | 19 | 1093 | | | | 100.0 | | | 0.23 | 103.64 | 80537 | 61.85 | 9567 | 0.31 | 0.07 | ↑ |
| 14 | 185 | 1067 | | | 100.0 | | | | 0.22 | 122.93 | 103982 | 80.51 | 17332 | 0.32 | 0.09 | ↑ |
| 15 | 2 | 1054 | | | | 100.0 | | | 0.22 | 101.35 | 64038 | 69.19 | 16220 | 0.35 | 0.10 | ↓ |
| 16 | 7 | 1032 | 9.0 | | 62.8 | 12.7 | 2.5 | 13.0 | 0.21 | 89.40 | 29828 | 65.52 | 24400 | 0.51 | 0.34 | ↑ |
| 17 | 99 | 991 | 100.0 | | | | | | 0.20 | 42.26 | 11646 | 30.61 | 7478 | 0.35 | 0.08 | ↓ |
| 18 | 18 | 947 | | 94.3 | 5.3 | | 0.4 | | 0.20 | 35.82 | 16928 | 33.60 | 9604 | 0.53 | 0.26 | ↓ |
| 19 | 3 | 875 | | | | | | 100.0 | 0.18 | 127.53 | 67426 | 30.08 | 7548 | 0.24 | 0.03 | ↑ |
| 20 | 44 | 862 | | 100.0 | | | | | 0.18 | 28.43 | 11916 | 8.08 | 7544 | 0.33 | 0.09 | ↓ |
| 21 | 175 | 858 | | | | | | 100.0 | 0.18 | 147.60 | 111692 | 86.77 | 10916 | 0.28 | 0.08 | ↓ |
| 22 | 13 | 838 | 100.0 | | | | | | 0.17 | 43.25 | 18197 | 28.42 | 6240 | 0.33 | 0.08 | ↓ |
| 23 | 181 | 807 | 83.5 | 16.0 | 0.4 | | | 0.1 | 0.17 | 86.69 | 84496 | 75.85 | 16102 | 0.36 | 0.28 | ↓ |
| 24 | 14 | 787 | | 99.7 | 0.1 | | | 0.1 | 0.16 | 106.77 | 73208 | 75.49 | 23445 | 0.35 | 0.12 | ↓ |
| 25 | 4 | 780 | | | | 100.0 | | | 0.16 | 101.71 | 24561 | 91.40 | 11309 | 0.30 | 0.06 | ↑ |
| 26 | 119 | 762 | | | | 100.0 | | | 0.16 | 89.81 | 58653 | 77.60 | 12990 | 0.28 | 0.06 | ↑ |
| 27 | 20 | 740 | | 100.0 | | | | | 0.15 | 30.18 | 12929 | 12.42 | 6104 | 0.26 | 0.13 | ↓ |
| 28 | 10 | 698 | 100.0 | | | | | | 0.14 | 21.32 | 3797 | 9.32 | 2233 | 0.28 | 0.07 | ↓ |
| 29 | 161 | 697 | | | 94.4 | 5.6 | | | 0.14 | 78.73 | 5204 | 77.77 | 3039 | 0.34 | 0.16 | ↑ |
| 30 | 60 | 632 | | | 100.0 | | | | 0.13 | 24.33 | 6462 | 13.54 | 3030 | 0.35 | 0.07 | ↓ |
| 31 | 43 | 607 | | 11.5 | 19.4 | 69.0 | | | 0.13 | 59.59 | 19260 | 49.40 | 19991 | 0.62 | 0.28 | ↓ |
| 32 | 87 | 606 | | 100.0 | | | | | 0.13 | 31.50 | 4585 | 24.89 | 2500 | 0.33 | 0.08 | ↓ |
| 33 | 23 | 600 | | | | | 100.0 | | 0.12 | 92.75 | 37891 | 55.18 | 20587 | 0.35 | 0.09 | ↓ |
| 34 | 108 | 599 | 0.2 | 84.0 | | 15.9 | | | 0.12 | 67.20 | 30105 | 71.38 | 30784 | 0.53 | 0.24 | ↓ |
| 35 | 57 | 597 | | 100.0 | | | | | 0.12 | 23.14 | 5942 | 6.87 | 3339 | 0.28 | 0.07 | ↓ |
| 36 | 458 | 596 | 73.5 | 0.2 | 3.7 | 0.2 | 16.1 | 6.4 | 0.12 | 60.54 | 51225 | 46.73 | 18196 | 0.57 | 0.41 | ↑ |
| 37 | 8 | 588 | 2.2 | | 1.4 | | | 96.4 | 0.12 | 105.11 | 71948 | 50.77 | 14150 | 0.47 | 0.19 | ↓ |
| 38 | 56 | 587 | | 100.0 | | | | | 0.12 | 31.57 | 12214 | 18.62 | 9787 | 0.55 | 0.24 | ↓ |
| 39 | 16 | 585 | | | | 99.8 | | 0.2 | 0.12 | 86.24 | 35435 | 55.66 | 10164 | 0.25 | 0.05 | ↑ |
| 40 | 33 | 584 | | 100.0 | | | | | 0.12 | 29.44 | 7644 | 12.18 | 2775 | 0.30 | 0.07 | ↓ |
| 41 | 15 | 584 | | 100.0 | | | | | 0.12 | 29.03 | 15927 | 9.09 | 6285 | 0.28 | 0.07 | ↓ |
| 42 | 463 | 583 | 78.4 | | 1.5 | 3.3 | 6.2 | 10.6 | 0.12 | 77.44 | 72906 | 53.72 | 27031 | 0.53 | 0.31 | ↑ |
| 43 | 231 | 578 | | | 0.5 | 99.5 | | | 0.12 | 177.20 | 113204 | 96.73 | 5841 | 0.28 | 0.08 | ↑ |
| 44 | 9 | 560 | 100.0 | | | | | | 0.12 | 131.64 | 87343 | 68.00 | 13501 | 0.33 | 0.06 | ↑ |
| 45 | 28 | 550 | | | 100.0 | | | | 0.11 | 38.64 | 17967 | 34.77 | 6896 | 0.27 | 0.07 | ↓ |
| 46 | 453 | 544 | 2.9 | 6.4 | 14.3 | | | 76.3 | 0.11 | 82.91 | 79073 | 67.98 | 15392 | 0.47 | 0.31 | ↑ |
| 47 | 17 | 544 | | | 100.0 | | | | 0.11 | 26.54 | 3363 | 7.93 | 1158 | 0.27 | 0.08 | ↓ |
| 48 | 177 | 539 | | | | 100.0 | | | 0.11 | 36.38 | 30000 | 25.17 | 11367 | 0.29 | 0.07 | ↓ |
| 49 | 21 | 533 | | | 99.6 | | | 0.4 | 0.11 | 78.57 | 20423 | 57.59 | 9723 | 0.32 | 0.11 | ↑ |
| 50 | 124 | 533 | | | | 100.0 | | | 0.11 | 79.38 | 43399 | 63.77 | 11638 | 0.32 | 0.09 | ↑ |

**Table S4.** Overview over the composition and scores of the ubiquitylation sites. All references to a score mean the ISA score. Clusters were selected by choosing clusters with the lowest mean ISA score and at least 90% composition of the respective ubiquitylation site.

| ubq site | total frames | frames in clusters | % clustered | % not clustered | µ ISA scores 4QLC | µ ISA scores 5NL0 | µ ISA scores 5WCU | µ ISA scores all | µ ISA scores off dyad | σ ISA scores 4QLC | σ ISA scores 5NL0 | σ ISA scores 5WCU | σ ISA scores all | σ ISA scores off dyad |
|---|---|---|---|---|---|---|---|---|---|---|---|---|---|---|
| K30 | 79398 | 28346 | 35.70 | 64.30 | 74.46 | 85.44 | 75.17 | 65.87 | 28.39 | 53.67 | 59.49 | 57.94 | 57.16 | 36.67 |
| K41 | 79398 | 23355 | 29.42 | 70.58 | 43.42 | 65.70 | 43.34 | 45.67 | 30.23 | 35.77 | 61.16 | 37.99 | 45.74 | 35.44 |
| K47 | 84198 | 26095 | 30.99 | 69.01 | 79.42 | 89.12 | 82.36 | 84.21 | 85.92 | 65.46 | 72.67 | 70.21 | 73.13 | 82.73 |
| K51 | 82193 | 41761 | 50.81 | 49.19 | 88.47 | 104.30 | 93.22 | 92.72 | 84.88 | 67.73 | 77.87 | 73.52 | 75.93 | 82.43 |
| K56 | 79398 | 7867 | 9.91 | 90.09 | 48.94 | 58.94 | 48.82 | 66.99 | 111.26 | 35.29 | 39.51 | 39.18 | 58.64 | 82.06 |
| K63 | 79398 | 15799 | 19.90 | 80.10 | 74.76 | 81.84 | 74.68 | 71.01 | 52.74 | 57.17 | 64.86 | 61.30 | 60.46 | 53.95 |

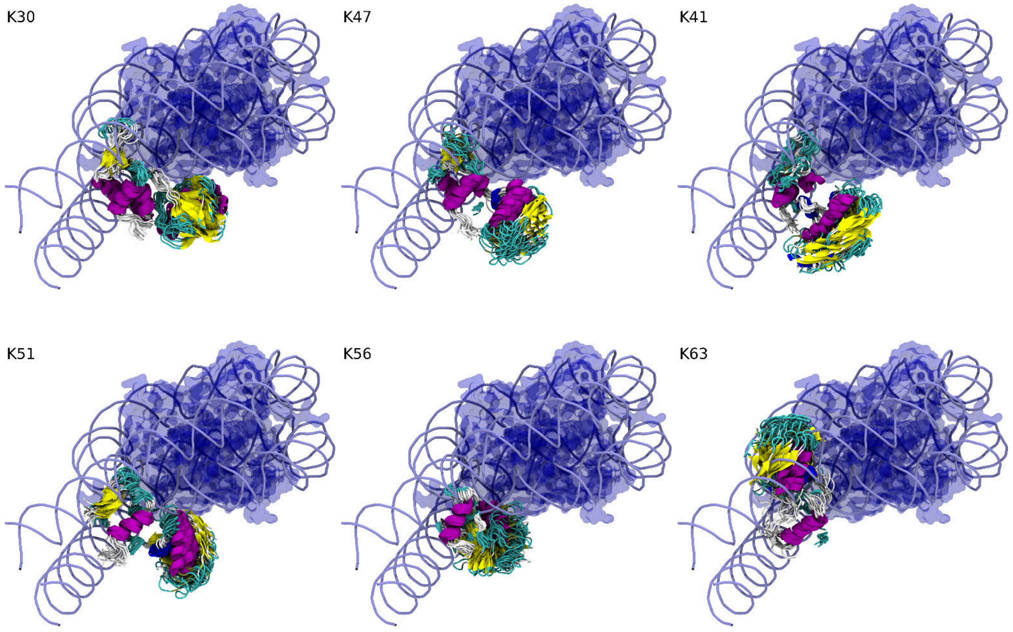


**Figure S10.** Best fitting clusters per ubiquitylation site using 5NL0 as the parent chromatosome. From all clusters found by iterative HDBSCAN clustering the cluster with the lowest ISA score containing at least 90% of a specific HUb variant is chosen. 20 equally spaced frames of these clusters are then placed into the parent 5NL0 chromatosome structure and visualized here. All of them have the Ub subunit pointing downward. For scores of other clusters refer to Table S3.

### S6 Manual Adjustments to linker histone positions in 1ZBB

The reference chromatosomes pdb 5NL0 and the off-dyad structure were manually adjusted to allow their linker histone to fit into the tetranucleosome array of pdb 1ZBB. In the following table these adjustments are listed. "Core" refers to one of the cores of 1ZBB, where core1 has the lower segid of the first frame of the biological assembly and core4 has the higher segid of the second frame of the biological assembly.

**Table S5.** Manual adjustments to create hypothetical linker histone positions in the tetranucleosome array 1ZBB.

| Alignment | origin | Adjustments made |
|---|---|---|
| 5NL0 to core 1 of 1ZBB | 1ZBB | rotated 5NL0 5 deg around third principal axis |
| 5NL0 to core 2 of 1ZBB | 1ZBB | rotated 5NL0 31 deg around third principal axis |
| 5NL0 to core 3 of 1ZBB | 1ZBB_frame_2 | rotated 5NL0 -5 deg around third principal axis |
| 5NL0 to core 4 of 1ZBB | 1ZBB_frame_2 | rotated 5NL0 -30 deg around first principal axis |
| off-dyad to core 1 of 1ZBB | 1ZBB | No additional adjustemts made |
| off-dyad to core 2 of 1ZBB | 1ZBB | -15 deg around third principal axis |
| off-dyad to core 3 of 1ZBB | 1ZBB_frame_2 | No additional adjustment made |
| off-dyad to core 4 of 1ZBB | 1ZBB_frame_2 | 8 deg around third principal axis |

#### References


1. Vijay-Kumar S, Bugg CE, Cook WJ. Structure of ubiquitin refined at 1.8 Åresolution. Journal of molecular biology. 1987;194(3):531–544.

2. Cerf C, Lippens G, Muyldermans S, Segers A, Ramakrishnan V, Wodak SJ, et al. Homo-and heteronuclear two-dimensional NMR studies of the globular domain of histone H1: sequential assignment and secondary structure. Biochemistry. 1993;32(42):11345–11351.

3. Pettersen EF, Goddard TD, Huang CC, Couch GS, Greenblatt DM, Meng EC, et al. UCSF Chimera—a visualization system for exploratory research and analysis. Journal of computational chemistry. 2004;25(13):1605–1612.

4. Van Der Spoel D, Lindahl E, Hess B, Groenhof G, Mark AE, Berendsen HJ. GROMACS: fast, flexible, and free. Journal of computational chemistry. 2005;26(16):1701–1718.

5. Schmid N, Eichenberger A, Choutko A, Riniker S, Winger M, Mark A, et al. testing of the GROMOS force-field versions: 54A7 and 54B7 Eur. Biophys J. 2011;40:843–856.

6. Berendsen HJ, Postma JP, van Gunsteren WF, Hermans J. Interaction models for water in relation to protein hydration. In: Intermolecular forces. Springer; 1981. p. 331–342.

7. Fincham D. Leapfrog rotational algorithms. Molecular Simulation. 1992;8(3-5):165–178.

8. Bussi G, Donadio D, Parrinello M. Canonical sampling through velocity rescaling. The Journal of chemical physics. 2007;126(1):014101.

9. Berendsen HJ, Postma Jv, van Gunsteren WF, DiNola A, Haak JR. Molecular dynamics with coupling to an external bath. The Journal of chemical physics. 1984;81(8):3684–3690.

10. Hess B, Bekker H, Berendsen HJ, Fraaije JG. LINCS: a linear constraint solver for molecular simulations. Journal of computational chemistry. 1997;18(12):1463–1472.

11. Darden T, York D, Pedersen L. Particle mesh Ewald: An N· log (N) method for Ewald sums in large systems. The Journal of chemical physics. 1993;98(12):10089–10092.

12. Berg A, Kukharenko O, Scheffner M, Peter C. Towards a molecular basis of ubiquitin signaling: A dual-scale simulation study of ubiquitin dimers. PLOS Computational Biology. 2018;14(11):1–14. doi:10.1371/journal.pcbi.1006589.

13. Ceriotti M, Tribello GA, Parrinello M. Simplifying the representation of complex free-energy landscapes using sketch-map. Proceedings of the National Academy of Sciences. 2011;108(32):13023–13028.

14. Leaver-Fay A, Tyka M, Lewis SM, Lange OF, Thompson J, Jacak R, et al. ROSETTA3: an object-oriented software suite for the simulation and design of macromolecules. In: Methods in enzymology. vol. 487. Elsevier; 2011. p. 545–574.
